# Phosphorylation of the exocyst subunit Exo70B2 contributes to the regulation of its function

**DOI:** 10.1101/266171

**Authors:** Ooi-Kock Teh, Chil-Woo Lee, Franck Anicet Ditengou, Till Klecker, Giulia Furlan, Marco Zietz, Gerd Hause, Lennart Eschen-Lippol, Wolfgang Hoehenwarter, Justin Lee, Thomas Ott, Marco Trujillo

## Abstract

The exocyst is a conserved hetero-octameric complex that mediates early tethering of post-Golgi vesicles during exocytosis. Its Exo70 subunit functions as a spatiotemporal regulator by mediating numerous interactions with proteins and lipids. However, a molecular understanding of the exocyst functions remains challenging. Exo70B2 localized to dynamic foci at the plasma membrane and transited through Brefeldin A (BFA)-sensitive compartments, indicating that it participates in conventional secretion. Conversely, treatment with the immunogenic peptide flg22 or the salicylic acid (SA) defence hormone analogue Benzothiadiazole (BTH), induced Exo70B2 transport into the vacuole where it colocalized with autophagic markers AUTOPHAGY-RELATED PROTEIN 8 (ATG8) and NEIGHBOR OF BRCA1 GENE 1 (NBR1). According with its role in immunity, we discovered that Exo70B2 interacts with and is phosphorylated by the MITOGEN-ACTIVATED PROTEIN KINASE 3 (MPK3). Mimicking phosphorylation inhibited Exo70B2 localization at sites of active secretion. By contrast, lines expressing phosphonull variants displayed higher Effector-Triggered Immunity and were hypersensitive to BTH, conditions known to induce the secretory pathway. Our results suggest a molecular mechanism by which phosphorylation of Exo70B2 regulates interaction with the plasma membrane, and couples the secretory pathway with cellular signalling.

## Introduction

The exocyst is an evolutionary conserved protein complex implicated in the tethering of secretory vesicles to the plasma membrane (PM) before soluble N-ethylmaleimide-sensitive-factor attachment receptor (SNARE)-mediated membrane fusion. The first exocyst components were described in yeast as temperature sensitive mutants in which vesicles accumulated due to failed fusion to the PM (Novick et al., 1980). All subunits of the exocyst were later shown to work in a complex and to mediate exocytosis of post-Golgi vesicles (Wu and Guo, 2015). The complex was subsequently found to be conserved in plants, and the first reported mutant phenotypes supported the anticipated function in secretion (Cole et al., 2005; Wen et al., 2005). However, in contrast to all other eukaryotic organisms, the Exo70 subunit has experienced a substantial expansion into many paralogues; 23 in *Arabidopsis thaliana* (referred to as Arabidopsis) and 47 in rice (Cvrckova et al., 2012).

The secretory pathway mediates the transport of lipids and proteins to the PM. Specific conditions, such as active growth and infection, engage the secretory pathway and significantly enhance its activity (Wang et al., 2005). During the immune response de-novo synthesis of a vast array of proteins is induced, including PM-localized receptors and proteins with antimicrobial activities which are delivered to the extracellular space (Gu et al., 2017). The defence hormone salicylic acid (SA) plays a key function by coordinately upregulating the secretory pathway (Wang et al., 2005; Nagashima et al., 2014).

At early stages of post-Golgi trafficking STOMATAL CYTOKINESIS DEFECTIVE1 (SCD1) and SCD2 proteins are though to recruit RabE1 and the exocyst to vesicles (Mayers et al., 2017). During the delivery of secretory vesicles, Sec3 and Exo70 subunits of the exocyst mediate binding to the PM (Ahmed et al., 2018). In plants, the ROP/RAC effector INTERACTOR OF CONSTITUTIVE ACTIVE ROPs 1 (ICR1) was shown to influence Sec3 localization (Lavy et al., 2007; Hazak et al., 2010). Localization of Exo70 in humans and yeast to the PM is mediated by its ability to bind phosphatidylinositol 4,5-bisphosphate (PI(4,5)P_2_) (He et al., 2007a; Liu et al., 2007). Although direct membrane binding of an exocyst component has not been shown in plants, the small molecule endosidin2 (ES2) binds and inhibits Exo70A1 exocytosis and recycling (Zhang et al., 2016; Mayers et al., 2017). Residues contributing to ES2 binding are located in the C-domain of Exo70A1 at its C-terminus, suggesting that the membrane-binding function of this region is conserved in plants.

The exocyst has been shown to play a role in the immune response of both plants and animals. The Sec5 subunit forms an effector complex with GTPase RalB and directly recruits and activates the atypical lκB kinase family member TBK1 in human cells (Chien et al., 2006). The RalB/Sec5 complex contributes to the activation of the innate immune response via TBK1 in response to virus infection. In plants, a function in immunity has mainly been reported for Exo70 homologues. Mutants of *Exo70B2* and *Exo70H1* are more susceptible to virulent bacterial pathogens (Pecenkova et al., 2011; Stegmann et al., 2012). Moreover, Exo70B2 was shown to be required for full activation of pathogen-associated molecular pattern (PAMP)-triggered immunity (PTI) (Stegmann et al., 2012). For Exo70B1, the closest homologue of Exo70B2, there have been contrasting reports regarding its function in the resistance against the bacterial pathogen *Pseudomonas syringae* pv. *tomato* DC3000 (Stegmann et al., 2012; Stegmann et al., 2013; Zhao et al., 2015).

Although the canonical function of the exocyst is exocytosis, it has been implicated in other processes (Wu and Guo, 2015). Exo70B1 is transported into the vacuole and partially colocalizes with AUTOPHAGY-RELATED PROTEIN 8 (ATG8), a ubiquitin-like protein required for the formation of autophagosomal membranes (Kulich et al., 2013). The Exo70E2 homologue is recruited to autophagosomes after autophagy induction by benzothiadiazole (BTH) (Lin et al., 2015). However, under control conditions, Exo70E2 remained in compartments that were distinct from autophagosomes. In human cells, RalB directly binds to Exo84, inducing the assembly of catalytically active ULK1 and Beclin1-VPS34 complexes on the exocyst during the activation of autophagosome formation (Bodemann et al., 2011).

A possible explanation for the various functions of the exocyst is that many cellular processes need to be coordinated with exocyst-mediated exocytosis (Wu and Guo, 2015). Therefore, exocyst activity must be tightly coordinated to allow temporal and spatial control of its functions.

In mammals, the mitogen-activated protein kinase ERK1/2 phosphorylates Exo70 in response to EPIDERMAL GROWTH FACTOR (EGF), resulting in the assembly of the exocyst complex and secretion (Ren and Guo, 2012). Adding another regulatory layer, we previously showed that upon activation of PTI in Arabidopsis, Exo70B2 was targeted by the E3 ligase PLANT U-BOX 22 (PUB22) for degradation during the immune response (Stegmann et al., 2012; Furlan et al., 2017).

In this study, we show that Exo70B2 can associate with the PM, and in contrast to other exocyst subunits, transits through a BFA-sensitive compartment. Treatment with the SA analogue BTH induces the accumulation of Exo70B2 in the microsomal membrane fraction, which reflects its transport into the vacuole. Exo70B2 can be phosphorylated by the PAMP-responsive MITOGEN-ACTIVATED PROTEIN KINASE 3 (MPK3), and plants expressing a non-phosphorylatable version display mislocalization of the protein at sites of active secretion. In addition, plants expressing a non-phosphorylatable version of Exo70B2 were more resistant to avirulent bacteria and more sensitive to BTH treatment. Together, our results reveal a close interplay between immune signalling, the secretory machinery and autophagy, in which Exo70B2 may act as a molecular rheostat to modulate secretory activity.

## Results

### Exo70B2 localizes to dynamic foci at the plasma membrane

To gain insight into exocyst function, and in particular of the Exo70B2 subunit, we characterized stable transgenic lines carrying the GFP-Exo70B2 fusion under control of the *UBQ10* promoter in the *exo70B2-3* background (Grefen et al., 2010). We opted to employ the *UBQ10* promoter, because lines under the control of a 1.5kbp fragment upstream of the *Exo70B2* CDS didn’t lead to detectable expression, similarly to what was reported by Li et al. (Li et al., 2010).

GFP-Exo70B2 was detected in the cytoplasm of epidermal cells in cotyledon and roots, as well as a continuous cell-peripheral localization, which suggested PM localization (Figure 1A). To confirm PM localization, we plasmolysed cells using mannitol. GFP-Exo70B2 was detected at the periphery and in Hechtian strands of plasmolysed epidermal cells, suggesting that Exo70B2 localized to the PM (Figure 1B). In addition, we analysed the localization in root hairs which show tip growth, reasoning that GFP-Exo70B2 localization would be focused to regions of high secretory activity (Honkanen and Dolan, 2016). High-resolution microscopy revealed that GFP-Exo70B2 localized to distinct foci at the periphery of root hairs, and the number of foci increased towards the tip where secretory activity is highest (Figure 1C). The additional analysis by variable angle epifluorescence (VEA) Total Internal Reflection (TIRF) microscopy demonstrated that these foci are dynamic with a transient dwelling at the PM (Movie S1).

**Figure 1.**
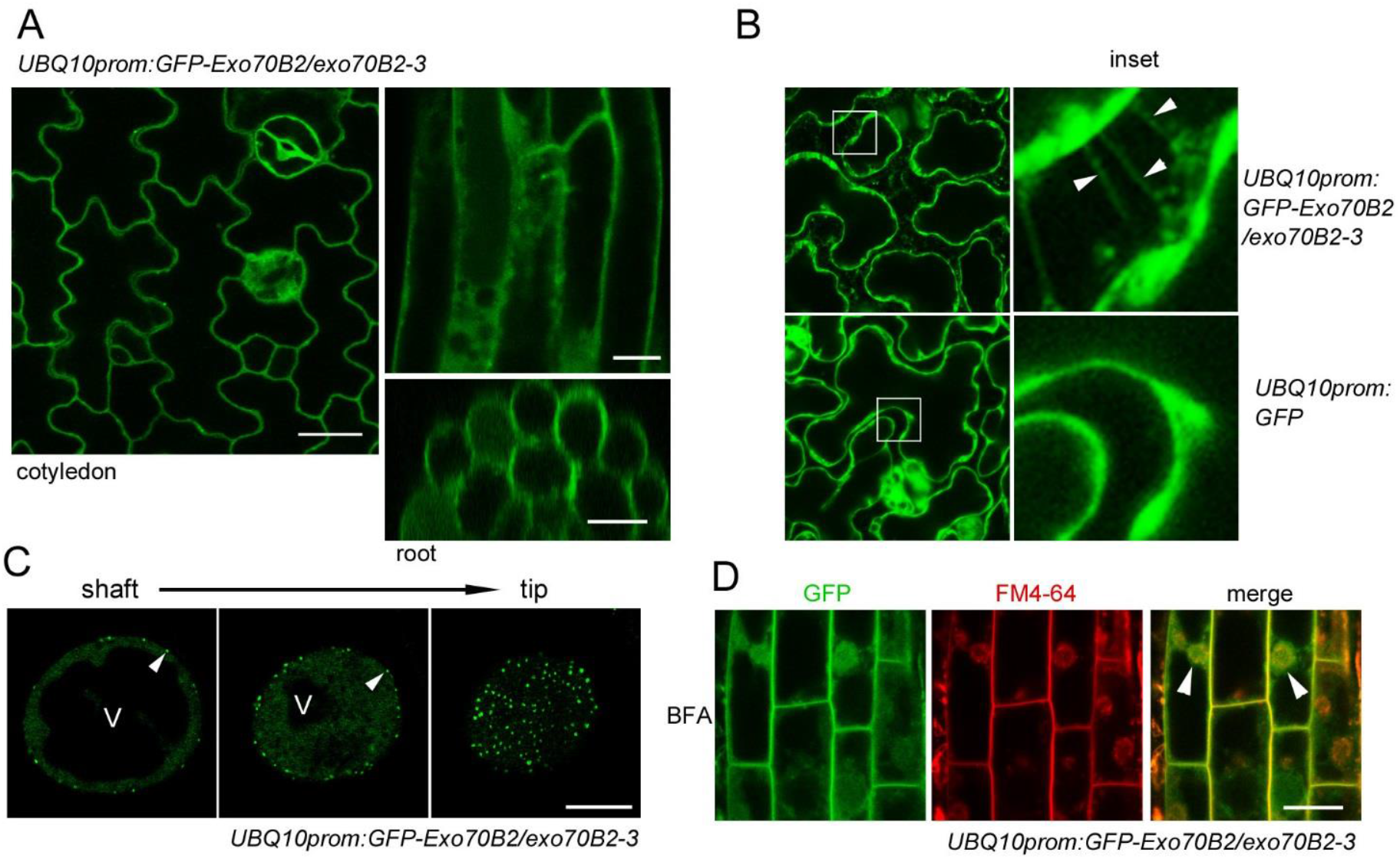
Exo70B2 localizes to the plasma membrane and transits through the trans-Golgi network. **(A)** Confocal laser-scanning microscopy images of four-day old homozygous transgenic seedlings expressing GFP-Exo70B2 under the control of the constitutive promoter *UBQ10* in the *exo70b2-3* background. Shown are epidermal cells of cotyledons (top left) and roots (right) and transverse section of root (right bottom). Size bars represent 20μm (cotyledons) and 10μm (roots). **(B)** *UBQWpro:GFP-Exo70B2/exo70b2-3* and *UBQ10pro:GFP* transgenic seedlings were incubated in 0.8M mannitol for 40 mins before analysed by CLSM. GFP-EX070B2 is visible on the Hechtian strands indicated by arrowheads. Scale bar 10μm.(C) High-resolution microscopy pictures of *UBQ10pro:GFP-Exo70B2/exo70b2-3* transgenic seedling root hairs. Three optical sections are shown from root hair shaft to tip. Scale bar 10μm, V denotes vacuole. **(D)** *UBQ10pro:GFP-Exo70B2/exo70b2-3* transgenic seedlings were stained with 5μM FM4-64 for 5 mins and subsequently incubated in 50μM BFAfor45 mins at room temperature. Scale bar 5μm.

Treatment with BFA, which inhibits endosomal recycling and secretion of PM proteins, resulted in the accumulation of Exo70B2 in BFA-bodies, indicating that it transits through the *trans*-Golgi network (TGN) (Figure 1D). These results support a function of Exo70B2 in conventional secretion of post-Golgi vesicles.

### Exo70B2 is transported into the vacuole upon activation of the immune response

We previously reported that Exo70B2 is targeted for degradation after activation of the immune response by the E3 ligase PUB22 (Stegmann et al., 2012). Ubiquitination of Exo70B2 by PUB22 suggested that it may be targeted for degradation via the proteasome. However, its association to the PM and accumulation in BFA-bodies indicated that Exo70B2 enters the endocytic, or alternate degradation pathways, capable of processing membrane-associated proteins.

To test this possibility, we treated seedlings with concanamycin A (ConcA), to inhibit vacuolar degradation, and flg22, which is perceived by the immune receptor FLAGELLIN-SENSITIVE 2 (FLS2) (Chinchilla et al., 2006), to activate the immune response. ConcA treatment alone did not result in the accumulation of Exo70B2 in the vacuole (Figure 2A). However, flg22/ConcA co-treatment revealed that Exo70B2 was recruited for vacuolar degradation (Figure 2A). As previously shown, flg22 treatment resulted in the degradation of Exo70B2 (Figure 2B).To test whether this reflected vacuolar degradation, we pre-treated seedlings with ConcA. Inhibition of the vacuolar acidification by ConcA reduced Exo70B2 proteolysis, hence supporting the notion that Exo70B2 is indeed degraded via the vacuole (Figure 2B).

**Figure 2.**
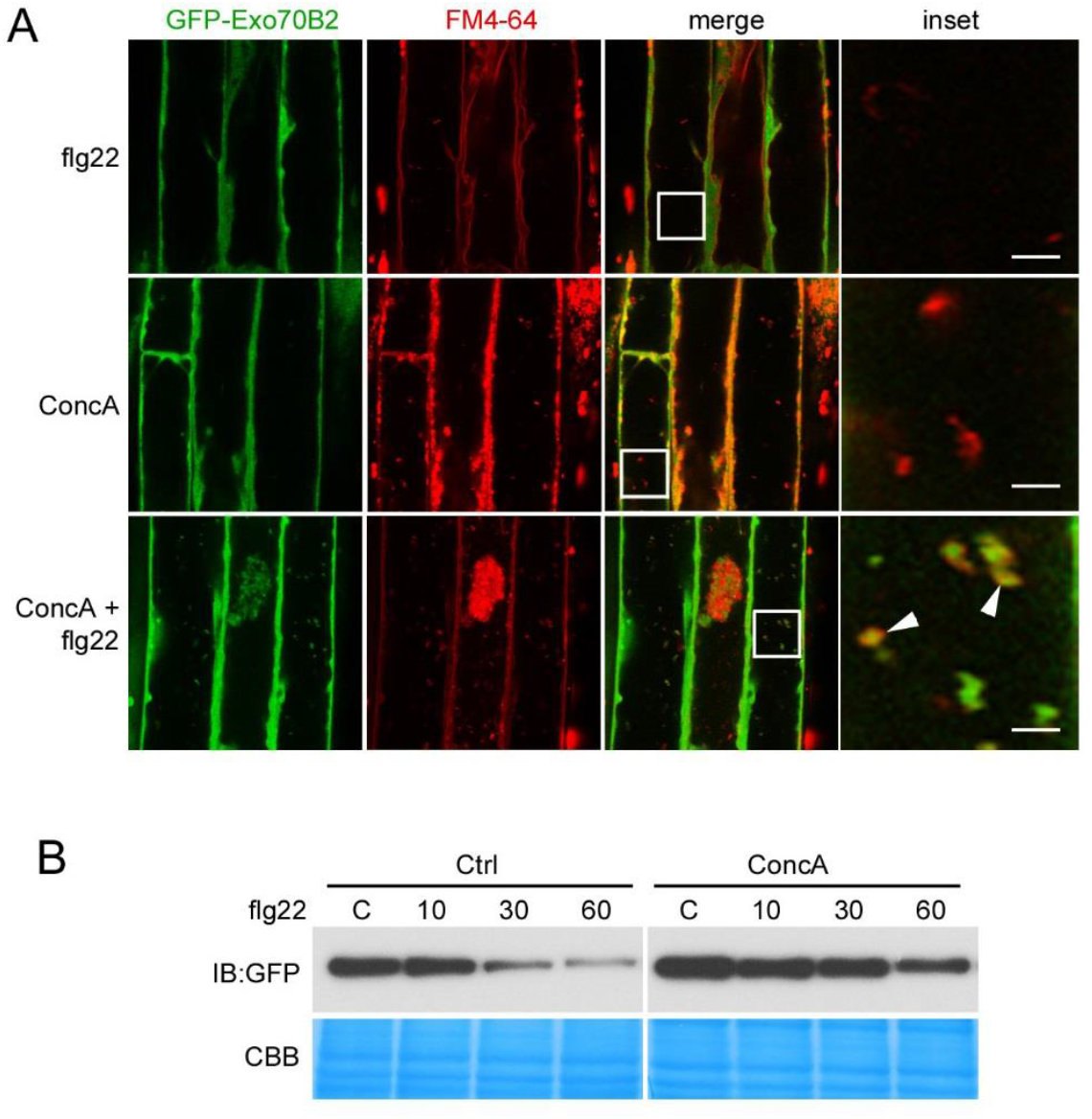
Exo70B2 is degraded during the immune response via the vacuole. **(A)** *UBQ10prom:GFP-Exo70B2/exo70b2-3* seedlings were treated with 1μM flg22 (1h), 1μM ConcA (2h) or 1μM flg22 (1h) + 1μM ConcA (2h) and subsequently stained for 5 mins in 5μM FM4-64. Arrowheads indicate vacuolar GFP-Exo70B2. Scale bar 5μm. **(B)** Two weeks old *UBQ10prom:GFP-Exo70B2/exo70b2-3* seedlings were treated with 1μM flg22 for the indicated times. Samples were mock treated (control) or 1μM ConcA was added 1h before elicitation. Total protein fractions were analysed by IB using the indicated antibody (shown are two cropped sections of the same blot). Equal loading is shown by CBB staining.

### Exo70B2 is transported into the vacuole where it colocalizes with autophagic markers

Previous studies in mammals have shown that the exocyst participates in the activation of autophagy (Bodemann et al., 2011). Moreover, Exo70B1 and Exo70E2 colocalized with the autophagy marker ATG8 upon tunicamycin or BTH treatment respectively, in plants (Ichimura et al., 2000; Kulich et al., 2013; Lin et al., 2015).

To induce autophagy we employed BTH (Yoshimoto et al.,2009; Lin et al., 2015), which is an analogue of the phytohormone SA, a central component of the immune response and induces the expression of hundreds of genes (Wang et al., 2005; Vlot et al., 2009). Inhibition of the vacuolar degradation by ConcA revealed basal transport of Exo70B2 into the vacuole, reflected by the accumulation of punctae in the vacuole of elongated epidermal root cells, but not detectable in cotyledon cells (Figure 3A). By contrast, co-treatment with BTH induced the accumulation of intravacuolar punctae in both roots and cotyledons (Figure 3A). Analysis of Sec6, another subunit of the exocyst, showed that BTH had a similar effect, and resulted in its delivery to the vacuole (Figure S1A).

**Figure 3.**
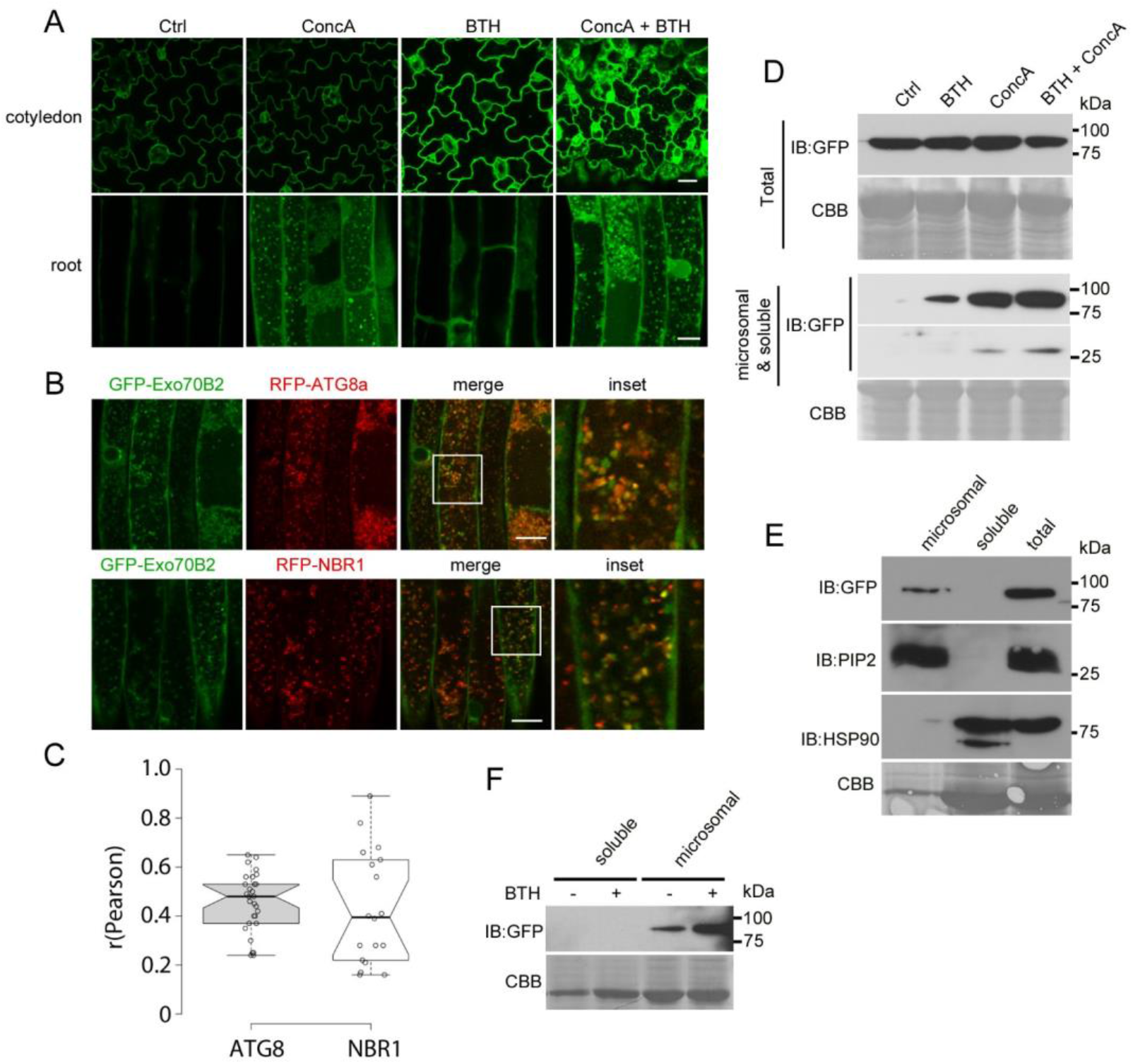
Exo70B2 is transported into the vacuole and colocalizes with autophagic markers. **(A)** GFP-EX070B2 is localised to autophagic body-like compartments after BTH and ConcA treatment. *UBQ10prom:GFP-Exo70B2/exo70b2-3* seedlings were treated o/n with either 0.1% DMSO, 1μM ConcA, 100μM BTH or 1μM ConcA + 100μM BTH. Confocal images of the same tissue were taken with identical imaging settings. Scale bars for cotyledon and root tissues are 20μm and 10μm, respectively. **(B)** GFP-Exo70B2 colocalizes with autophagosome markers ATG8a and NBR1. Double transgenic lines carrying *UBQ10prom:GFP-Exo70B2* and *UBQ10prom:RFP-ATG8a* or *UBQ10prom:RFP-NBR1* were treated overnight with 1μM ConcA + 100μM BTH before CLSM analysis. Scale bar 10μm. **(C)** Co-localisation analysis for GFP-Exo70B2 with mCherry-ATG8a or mcherry-NBR1. Data points represent the technical replicates of 2 independent experiments. Boxplots show median and inter quantile range (IQR), outliers (> 1.5 times IQR) are shown as circles. **(D)** *UBQ10pro:GFP-Exo70B2/exo70b2-3* transgenic seedlings were treated with 0.1% DMSO (control), 100μM BTH overnight (o/n), 1μM ConcA o/n and indicated combinations. Samples from the same experiment were aliquoted, and total protein (detergent-solubilized) or soluble and microsomal fraction (supernatant from centrifugation step 5′ at 12k ref) were resolved by PAGE and analysed by IB using anti-GFP antibodies (shown are two cropped sections of the same blot). Equal loading is shown by CBB staining. Similar results were obtained in three biological replicates. **(E)** Total protein, soluble and microsomal fractions from *UBQ10pro:GFP-Exo70B2/exo70b2-3* transgenic seedlings were resolved by PAGE and analysed by IB using the indicated antibodies. Shown are equivalent protein amounts from different fractions. Plasma membrane intrinsic protein2 (PIP2) and heat shock protein 90 (HSP90) were used as markers for microsomal and soluble fractions, respectively. **(F)** Soluble and microsomal fractions of *UBQ10pro:GFP-Exo70B2/exo70b2-3* seedlings were analysed by IB after 100μM BTH o/n treatment. Experiments in E and F were repeated with similar results.

Supporting the transport of Exo70B2 into the vacuole by autophagy, the marker proteins ATG8a and NBR1 colocalized to a subpopulation of punctae after BTH/ConcA treatment (Figure 3B) with Pearson’s correlation coefficients of 0.48 for ATG8 and 0.4 for NBR1 (Figure 3C) (Kirkin et al., 2009; Svenning et al., 2011). Moreover, ultrastructural analysis confirmed that Exo70B2 was transported into the vacuole upon autophagy induction (Figure S1B).

To further dissect the behaviour of Exo70B2, we analysed the effect of BTH on protein levels. Analysis of total protein detected by immunoblot (IB) did not show obvious changes after BTH treatment, in spite of increased fluorescence intensity (Figure 3D, upper panel). Similarly, ConcA treatment, alone or in combination with BTH, resulted in only minor variations of Exo70B2 in total protein extracts. Because BTH treatment remobilises Exo70B2 to the vacuole, we analysed protein fractions containing both soluble and microsomal proteins, but lacking larger organelles such as nuclei and chloroplasts. In contrast to total protein, this fraction contained non-detectable levels of Exo70B2 in control samples (Figure 3D, lower panel). However, treatment with BTH and ConcA, alone or combined, resulted in increased amounts of Exo70B2. Moreover, GFP cleavage, which can be indicative of autophagy (Marshall et al., 2015), was observed for samples treated with ConcA, BTH and ConcA (Figure 3D, lower panel).

This fraction was further separated by ultracentrifugation, resulting in a microsomal pellet containing membrane-bound proteins, and a supernatant with soluble proteins. Comparison of equivalent sample amounts showed that while Exo70B2 was not detectable as a soluble protein, it was present in the microsomal fraction (Figure 3E). Finally, we also show that the microsome-associated Exo70B2 increased upon BTH treatment (Figure 3F).

Since flg22 treatment induced Exo70B2 transport into the vacuole, we wondered whether flg22 potentially had an effect on autophagy. Treatment of seedlings with flg22 resulted in an increase of NBR1 protein levels, and flg22/ConcA co-treatment further enhanced this effect (Figure S2A). In accordance, NBR1 accumulated in the vacuole in response to flg22, and co-treatment with ConcA resulted in both a diffuse vacuolar signal and perivacuolar punctae (Figure S2B). These results are in agreement with a potential degradation of Exo70B2 by autophagy in response to the activation of the immune response.

Taken together, BTH activates Exo70B2 transport into the vacuole potentially by autophagy, which is reflected by the colocalization with ATG8 and the remobilization of Exo70B2 sub-pools to membrane fractions.

### Exo70B2 interacts with MPK3 and is phosphorylated in residues located in the C-domain

Induction of Exo70B2 transport into the vacuole by flg22 treatment prompted us to search for a link to immune signalling. Exocyst subunits including Exo70B2 were reported to be significantly increased in phosphoprotein-enriched fractions of Arabidopsis plants expressing the constitutively active MITOGEN-ACTIVATED PROTEIN KINASE KINASE 5 (MKK5)-DD variant in comparison to the kinase-inactive KR mutant: Exo70B2 by 2.28 fold (p value <0.001), Exo70E1 2.35 fold (p value <0.05) and Sec5A 2.86 fold (p value <0.05) (Lassowskat et al., 2014; Lee et al., 2015). FLS2 activation by flg22 triggers the MKK5 branch of MAPKs, which includes MPK3 and MPK6 and other downstream kinases. Because MAPK signalling is induced during the immune response, we hypothesized that Exo70B2 may be regulated by phosphorylation.

To evaluate this possibility, we first assayed the interaction with MPK3, MPK4, MPK6, MPK8 and MPK11 using bimolecular fluorescence complementation (BiFC). We found that from the tested MAPKs, both MPK3 and MPK11 reconstituted fluorescence when transiently coexpressed with Exo70B2 (Figure 4A and Figure S3A). Although the BiFC results suggested that Exo70B2 also interacts with MPK11, the lack of specific MPK11 antibodies and the typically low activity of MPK11 (Bethke et al., 2012), impedes its analysis. Therefore, we decided to focus on MPK3, which was also shown to phosphorylate the E3 ligase PUB22, which targets Exo70B2 (Furlan et al., 2017). Confirming the interaction, endogenous MPK3 co-immunoprecipitated (IP) with Exo70B2 by anti-GFP antibodies in two independent homozygous lines (T3) carrying the *UBQ10prom-GFP-Exo70B2* construct in the *exo70B2-3* background (Figure 4B). The interaction was independent of flg22 treatment, suggesting that both proteins are in complex under the tested conditions. In vitro pull-down experiments of bacterially expressed Exo70B2 and MPK3 showed that both proteins interacted directly, while MPK6 displayed only a weak interaction (Figure 4C).

**Figure 4.**
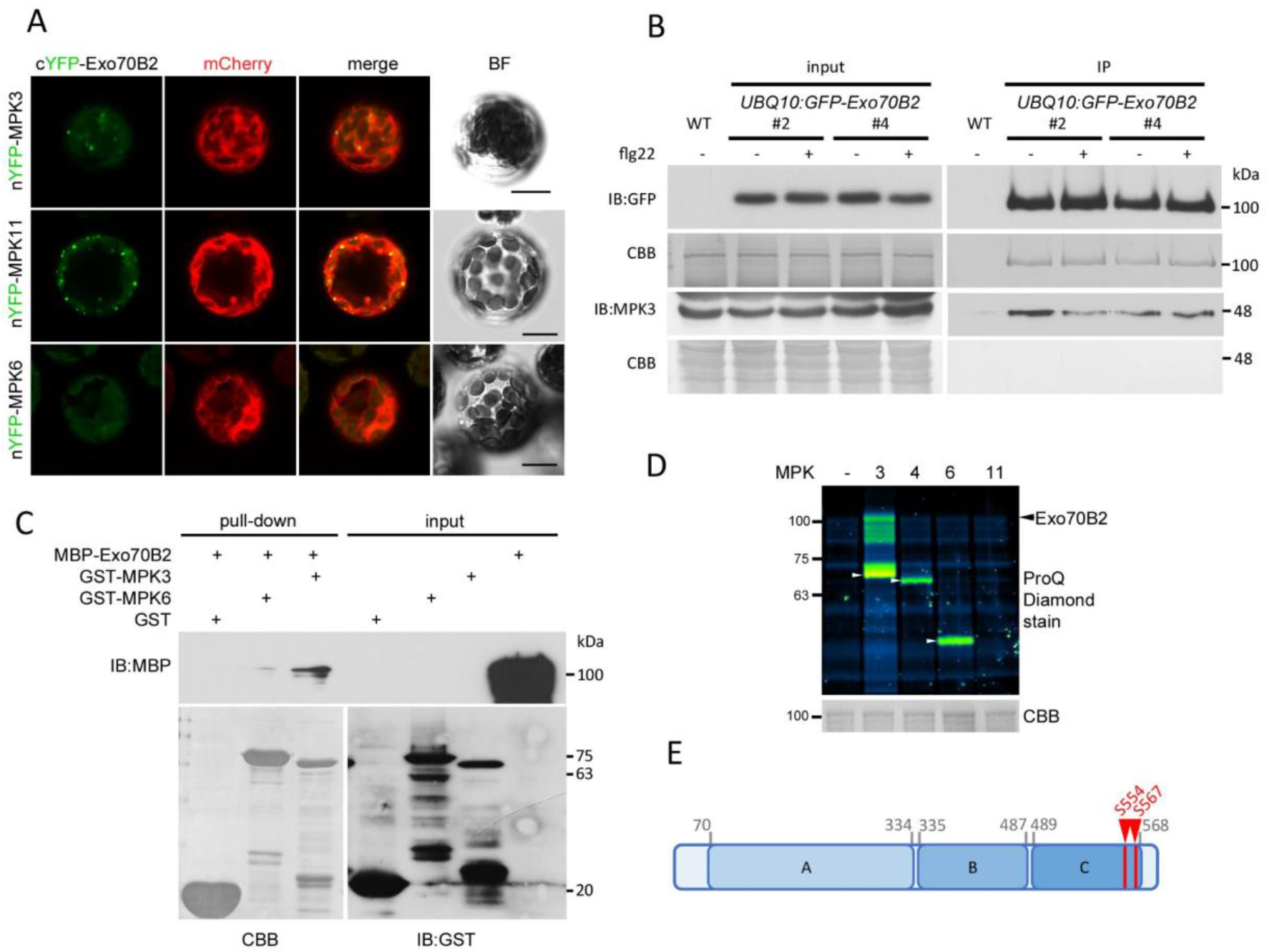
EX070B2 is phosphorylated at residues S554 and S567. **(A)** Interaction between Exo70B2 and MPK3 detected by BiFC in Arabidopsis protoplasts. nYFP-MPK3, nYFP-MPK6 or nYFP-MPK11 were coexpressed with cYFP-Exo70B2 as indicated. Free mCherry was coexpressed to label the cytoplasm and nucleus. Scale bar 50μm. Experiment was repeated with similar results. **(B)** Transgenic seedlings carrying *UBQ10prom:GFP-Exo70B2* were treated with 1μM flg22 for 20min and subjected to IP with anti-GFP beads. Endogenous coimmunoprecipitated MPK3 was detected with MPK3-antibodies. **(C)** MBP-Exo70B2 pull-down assay using bacterially expressed and purified GST-MPK3 and GST-MPK6 on glutathione agarose beads as baits. **(D)** GST-Exo70B2 was incubated alone or with activated GST-MPK3, GST-MPK4 and untagged MPK6 (white arrowheads). MPK11 could not be activated by MKK5. Phosphorylation was visualized with ProQ Diamond stain. White arrows indicate autophosphorylated kinases. **(E)** Cartoon depicting the localization of phosphorylated sites in Exo70B2 identified by LC-MS/MS from in vitro phosphorylation assays with GST-MPK3 and in vivo from GFP-Exo70B2 immunopurified from transgenic lines treated with flg22.

Interaction between Exo70B2 and MPK3 opened the possibility that Exo70B2 is a substrate of MPK3. To test this, we carried out in vitro phosphorylation assays using purified recombinant MPK3, MPK4, MPK6 and MPK11. Consistent with the direct interaction, Exo70B2 was readily trans-phosphorylated by MPK3 but not by MPK4 or MPK6 (Figure 4D). In the case of MPK11, kinase activity of recombinant MPK11 is typically low in our hands, and consequently, no phosphorylation of Exo70B2 was detectable.

Recombinant Exo70B2 phosphorylated by MPK3 was next analysed by LC-MS/MS to determine the phosphorylation sites. We identified two residues, namely S554 and S567, both of which are located in the C-domain at the C-terminal portion of the protein (Figure 4E, Figure S3B and Table S1). Both residues are followed by a proline, and thus represent typical MAPK phosphorylation motifs. No phosphopeptides were identified in controls without kinase. To confirm that the phosphorylation also takes place in vivo, we immunopurified GFP-Exo70B2 from transgenic seedlings treated with flg22 for 30 and 140 min and analysed samples by LC-MS/MS. Modifications in residues S554 and S567 could be identified, indicating that phosphorylation also takes place in vivo, while no phosphorylation was detected in non-treated samples (Figure S3C and Table S1). A sequence alignment between Exo70 homologues of the identified sites indicates that these residues are not conserved, suggesting a specialized regulatory function in Exo70B2 (Figure S3D).

### Phosphorylation of Exo70B2 regulates its localization at sites of active secretion

Together with Sec3, the Exo70 subunit mediates tethering of the exocyst by interacting with phosphoinositides at the PM via ionic interactions (He et al., 2007a; Liu et al., 2007). The identified phosphorylation sites in Exo70B2 are in the vicinity of the corresponding amino acids shown to affect membrane interaction in the mouse Exo70 (violet) (He et al., 2007a), as well as residues that contribute to ES2 binding in Exo70A (green, Figure 5A and Figure S3D) (Zhang et al., 2016), suggesting that they could affect Exo70B2 membrane binding. To test whether phosphorylation of S554 and S567 had an impact on the interaction on Exo70B2’s localization, we generated lines expressing a non-phosphorylatable S554/567A, as well as a phosphomimetic S554/567D variant. In order to detect potential effects of protein localization we again analysed root hairs by high resolution microscopy. Seedlings expressing the WT protein, displayed an even distribution of Exo70B2 punctae in root hair tips (Figure 5B). In plants expressing the S554/567A variant, Exo70B2 was localized to similar punctate compartments, however, it also accumulated in larger patches (Figure 5B). By contrast, the punctae were barely detectable for the phospho-mimetic S554/567D variant (Figure 5B). A structural model of Exo70B2’s C-domain suggests that phosphorylation sites are exposed and placed adjoining the last stretch of a basic ridge (Figure S4).

**Figure 5.**
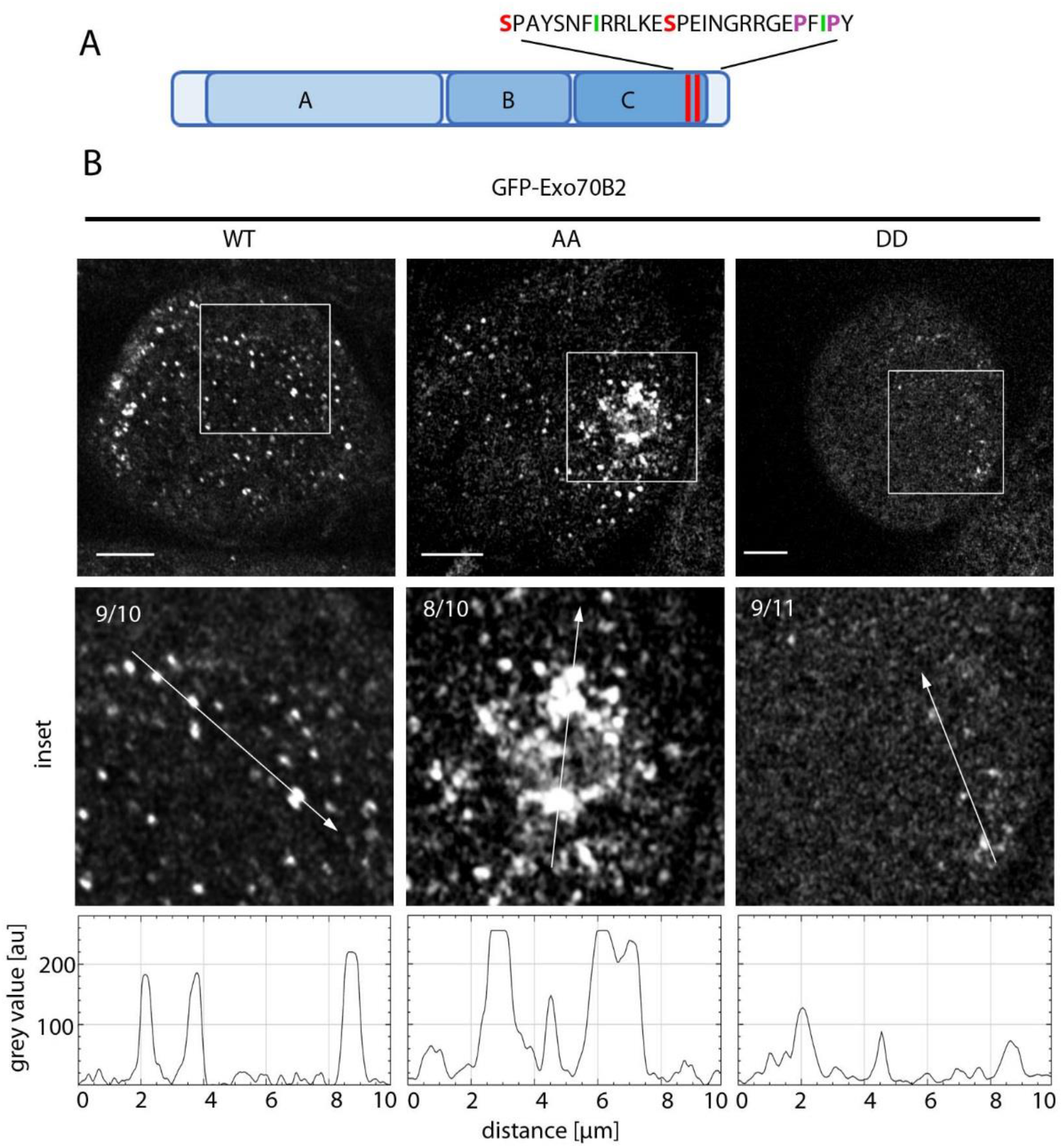
Ser554 and Ser567 regulate Exo70B2 localization in root hairs. **(A)** Cartoon depicting the identified phosphorylation sites Ser554 and Ser567 (red) in Exo70B2 C-domain, and corresponding amino acids contributing to ES2 binding in Exo70A1 (green) and membrane binding of the baker’s yeast Exo70 (purple). **(B)** High-resolution microscopy pictures of *UBQ10pro:GFP-Exo70B2/exo70b2-3* transgenic seedling root hairs expressing the WT, S554/567A (AA) or S554/567D (DD) proteins. Three optical sections taken from the cortical layer. Scale bar 10μm. Inset shows a magnification and the arrows denote pixel-intensity measurement shown below. Numbers show the ration of root hairs displaying the respective localization. Experiment was repeated with similar results.

### Exo70B2 interaction with ATG8 through AIMs is increased in phosphomimic variant

Sequence analyses have predicted an over proportional presence of ATG8-interaction motifs (AIMs) in the Arabidopsis Exo70 family, although it remains unknown whether these are functional (Cvrckova and Zarsky, 2013; Tzfadia and Galili, 2013). We identified two putative AIMs also located in the C-domain of Exo70B2: AIM1 508-DGP<u>Y</u>PK<u>L</u> and AIM2 522-SQ<u>F</u>DE<u>V</u> (Figure 6A).

**Figure 6.**
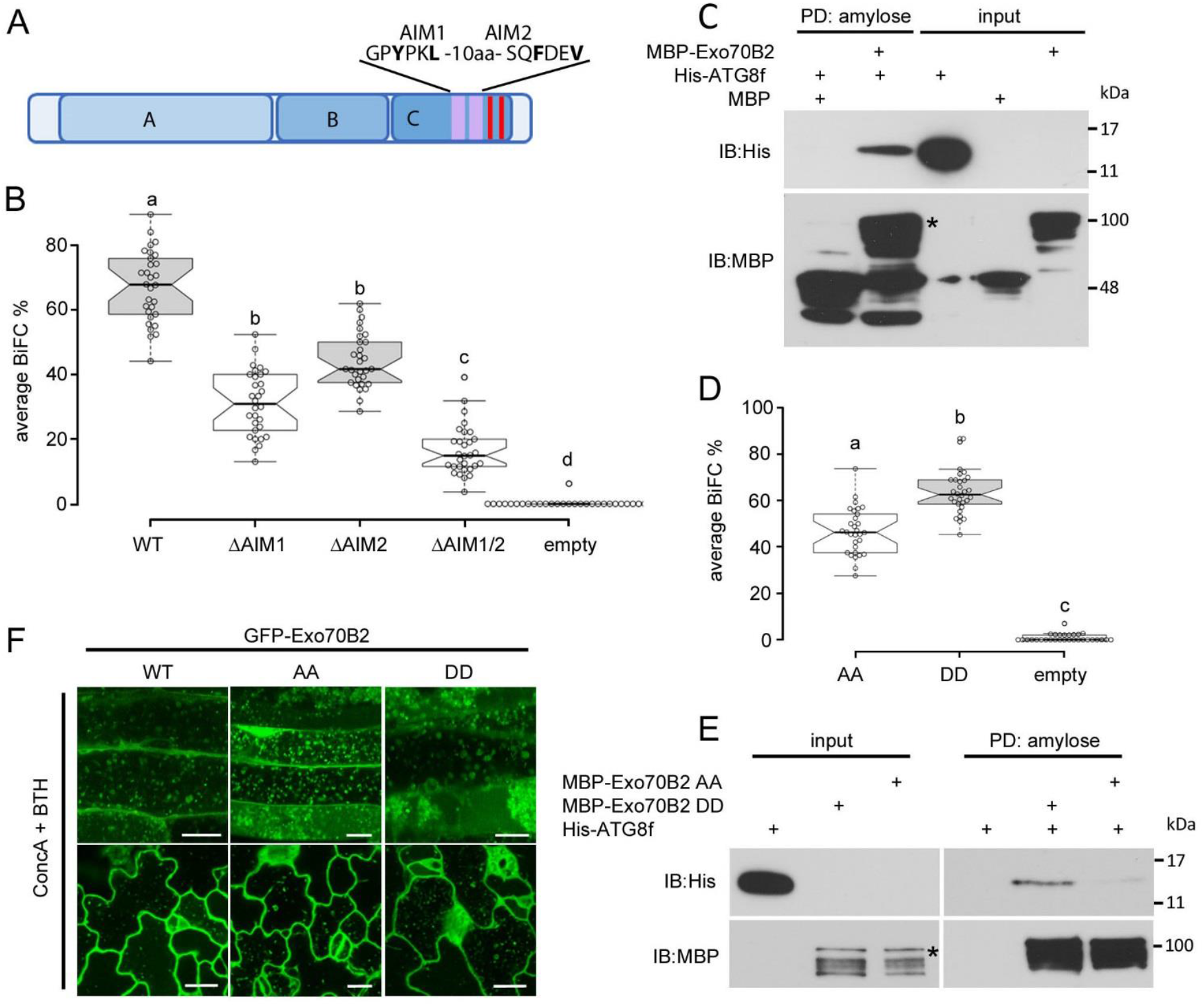
Exo70B2 interacts with ATG8 via C-terminal AIMs. **(A)** Cartoon depicting the localization and sequences of ATG8-interacting motifs (AIMs) highlighted in violet. Phosphorylation sites are highlighted in red. **(B)** BiFC of coexpressed nYFP-ATG8f and either cYFP-Exo70B2 (WT) or mutant variants cYFP-Exo70B2-ΔAIM1 (ΔAIM1), cYFP-Exo70B2-ΔAIM2 (ΔAIM2), cYFP-Exo70B2-ΔAIM1/AIM2 (ΔAIM1/2) or empty pSPYCE. Constructs were transiently coexpressed in Arabidopsis mesophyll protoplasts. mCherry was transformed as a marker for transformation. Boxplots show median and inter quantile range (IQR), outliers (> 1.5 times IQR) are shown as circles. Letters indicate statistically significant differences (One-Way ANOVA and Tukey post hoc test, p<0.05). Percentages of fluorescence complementation obtained from 30 independent images, each with 25-40 transformed protoplasts. Total protoplasts scored for WT, ΔAIM1, ΔAIM2, ΔAIM1/2 and empty vector were 723, 754, 708, 683 and 542, respectively. The experiment was repeated with similar results. **(C)** Pull-down (PD) assay with bacterially expressed and purified recombinant proteins. His-ATG8f was co-purified using MBP-Exo70B2 on amylose agarose beads. Asterisk indicates MBP-Exo70B2. **(D)** BiFC of coexpressed nYFP-ATG8f and either cYFP-Exo70B2 S554/567A (AA), S554/567D (DD) or empty vector were transiently coexpressed in Arabidopsis mesophyll protoplasts. mCherry was used as a transformation marker. Evaluation of percentage of fluorescence complementation and statistical analysis as in (B). Total protoplasts scored for Exo70B2 WT, AA, DD and empty vector were 1283, 695, 1325 and 1302, respectively. The experiment was repeated with similar results. **(E)** Pull-down assay using recombinant His-ATG8f and MBP-Exo70B2 S554/567A (AA) or S554/567D (DD) on amylose agarose beads. Asterisk indicates MBP-Exo70B2. **(F)** Transgenic seedlings expressing GFP-Exo70B2 WT, S554/567A (AA) and S554/567D (DD) were treated overnight in 1μM ConcA + 100μM BTH before CLSM analysis. Scale bars are 20μm for cotyledon and 10μm for root tissues.

We first tested the interaction by coexpressing Exo70B2 with ATG8f, which resulted in fluorescence complementation in a BiFC assay (Figure 6B and Figure S5A). To confirm that the interaction was mediated by the AIMs, we generated mutants in which we replaced <u>Y</u>PK<u>L</u> (AIM1) and <u>F</u>DE<u>V</u> (AIM2) by <u>A</u>PK<u>A</u> and <u>A</u>DE<u>A</u>, respectively. Mutation of single AIMs significantly compromised complementation in comparison to WT Exo70B2, and was mostly abrogated in the ΔAIM1/AIM2 double mutant (Figure 6B). To further characterize the interaction, we carried out in vitro pull-down assays with purified recombinant proteins. MBP-Exo70B2 was able to co-purify His-ATG8f, indicating that the proteins interact directly (Figure 6C). In contrast to AIM1, which is located in a highly variable region of the Arabidopsis Exo70 family, AIM2 is present in 65% of the Arabidopsis Exo70 homologues, and conserved in *Saccharomyces cerevisiae* Exo70 (Figure S5B).The conservation of the second AIM in the yeast Exo70 opened the possibility that the interaction with ATG8 and its transport into the vacuole represent an ancestral function. We therefore, generated a yeast strain expressing an Exo70-FLAG fusion protein from its genomic locus and transformed these cells with a plasmid carrying GFP-ATG8. Under nutrient replete conditions Exo70 interacted with ATG8 (Figure S5C), and the interaction was markedly increased after 1h nitrogen (N) deprivation (Figure S5C). These results show that the interaction between Exo70 and ATG8 is conserved across kingdoms and uncover a direct link to autophagy.

The identified phosphorylation sites are located in the vicinity of the AIMs, suggesting a potential role in their regulation (Figure 6A and Figure S4). We tested whether phosphorylation had an impact on the interaction with ATG8f. Coexpression of ATG8f with Exo70B2 S554/567A displayed a significantly decreased YFP complementation, when compared to S554/567D (Figure 6D and Figure S5D), suggesting that phosphorylation of serines increases Exo70B2’s ability to interact with ATG8f. Furthermore, in vitro pull-down assays show that in comparison to the Exo70B2 S554/567A, ATG8f co-precipitated more efficiently by the S554/567D phosphomimetic variant, suggesting a stronger interaction (Figure 6E).

Since Exo70B2 may be transported into the vacuole by autophagy, we tested a potential role of S554 and S567 phosphorylation. However, although the phosphomimetic interacted better with ATG8f, both mutant variants were still delivered to the vacuole under the tested conditions (Figure 6F). Together, we show that Exo70B2 contains two functional AIM motifs, and that phosphorylation at S554 and S567 may control ATG8f interaction but is dispensable for its transport into the vacuole.

### Expression of Exo70B2 phosphonull variant enhances sensitivity to BTH and resistance to avirulent bacteria

Because secretion is induced during the immune response and is key to mount and deploy immune responses (Gu et al., 2017), we carried out pathogen infection assays using the virulent *Pseudomonas syringae* pv. *tomato* DC3000 (*Pst*), as well as an avirulent *Pst* strain expressing the AvrRPS4 effector, which is recognized in Arabidopsis and activates effector-triggered immunity (ETI). As previously reported, *exo70B2-3* was more susceptible to the virulent *Pst* when compared to the WT Col-0 (Figure 7A). Re-introducing the *GFP-Exo70B2* complemented the enhanced susceptibility back to WT levels (Figure 7A). The S554/567A and S554/567D GFP-Exo70B2 mutants, however, were also able to complement the phenotype to a similar extent as the WT protein. By contrast, during ETI, while no significant differences were observed for most lines, plants expressing the phosphonull S554/567A variant, were significantly more resistant to *Pst-AvrRPS4* (Figure 7B).

**Figure 7.**
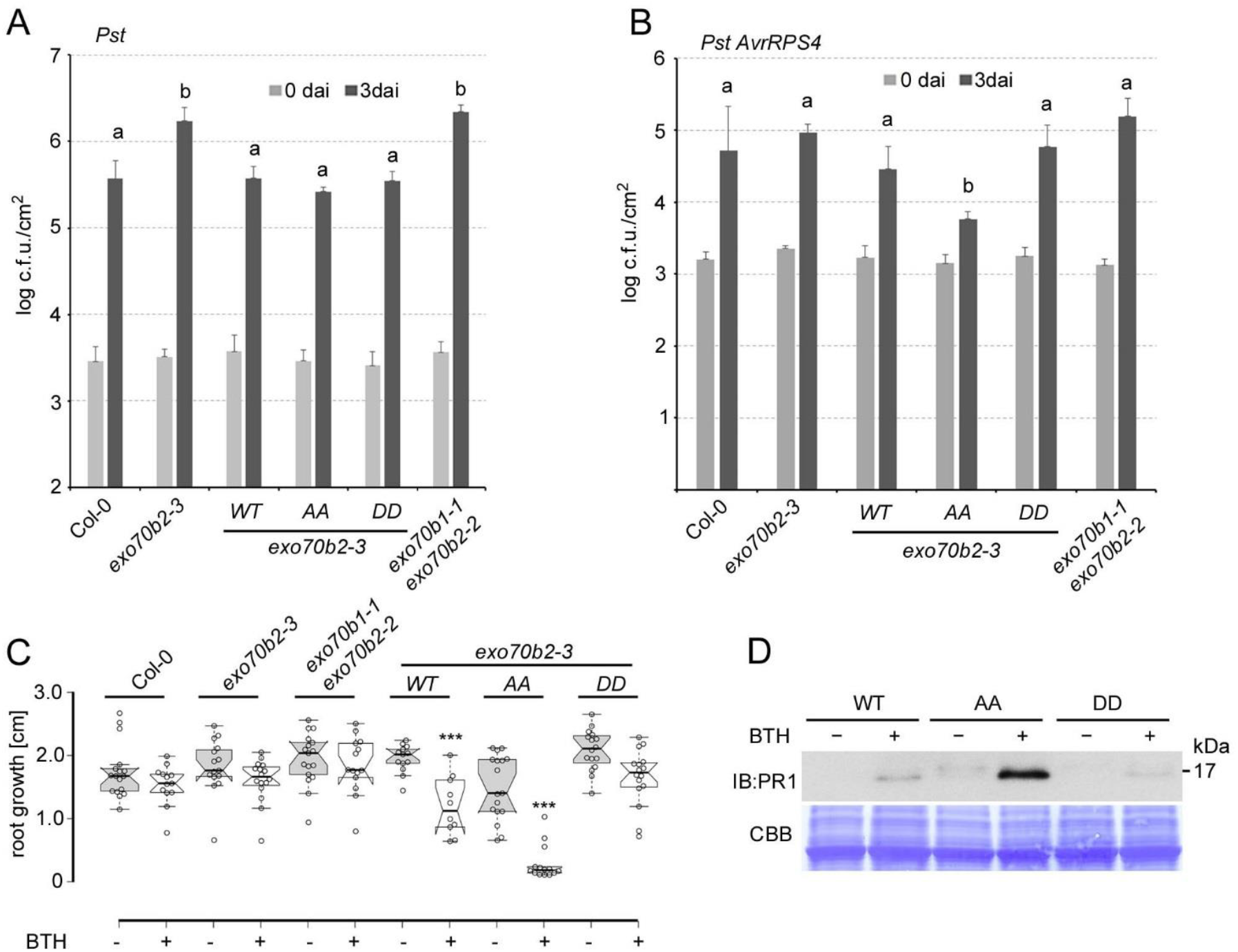
Plants expressing Exo70B2 phosphonull variant are more sensitive to BTH and resistant to avirulent bacteria. **(A)** Infection assays with the virulent bacterial pathogen *Pseudomonos syringae* pv. *tomato* DC3000 (*Pst*) empty vector. Six week-old plants were spray inoculated with a bacterial suspension of 5×10^8^ c.f.u./mL and analysed 0 and 3 days after inoculation. Data shown as mean +/− S.D. (n = 5). Letters indicate statistically significant differences between c.f.u. in different lines three days after inoculation (dai, One-Way ANOVA and Tukey post hoc test, p < 0.05). Similar results were obtained in three independent experiments. **(B)** Infection assays with the avirulent bacterial pathogen *Pseudomonas syringae* pv. *tomato* DC3000 *AvrRPS4*. Six week-old plants were syringe-infiltrated with a bacterial suspension O.D._600_ 0.001 and analysed 0 and 3 days after inoculation. Data shown as mean +/− S.D. (n = 5). Letters indicate statistically significant differences (One-Way ANOVA and Tukey post hoc test, p < 0.05). **(C)** Primary root lengths of *UBQ10pro:GFP-Exo70B2/exo70b2-3* transgenic seedlings expressing GFP-Exo70B2 WT, S554/567A (AA), S554/567D (DD) and ΔC-domain (ΔC) measured seven days after transplanting onto media +/− 100μM BTH. Boxplots show median and IQR, outliers (> 1.5 times IQR) are shown as circles. Values of a representative experiment. Asterisks indicate statistically significant differences between control and BTH treatment (One-Way ANOVA and Tukey post hoc test, p < 0.05). Experiment was repeated four times with similar results. See also Table S2 for complete statistical analysis results. **(D)** Transgenic seedlings described in (A) were treated with 100μM BTH overnight (o/n). Total protein samples were resolved by PAGE and analysed by IB using anti-PR1 antibodies. Equal loading is shown by CBB staining. The experiment was repeated with similar results.

During ETI, plants trigger the hypersensitive response, which is a robust immune reaction characterized by programmed cell-death and the concomitant accumulation of high levels of SA (Jones and Dangl, 2006). Similar to SA, BTH induces the coordinated up-regulation of the secretory pathway (Wang et al. 2003). We therefore, investigated the impact of Exo70B2 phosphorylation by scoring seedling root growth in the presence of BTH. BTH had little to no effect on root growth of Col-0 plants when compared with controls under the used conditions (Figure 7C and Table S2). Similarly, no major changes were observed for the other lines, with the exception of *exo70B2-3* plants complemented with WT Exo70B2 and the S554/567A phosphonull variant, which showed root growth inhibition in the presence of BTH (Figure 7C and Table S2). Of note, the S554/567A phosphonull mutant was significantly more sensitive than WT Exo70B2 expressing lines, displaying growth arrest after transplanting them onto BTH-containing media. By contrast, the S554/567D phosphomimetic variant showed no significant difference to the control (Figure 7A and Table S2). To further characterize the BTH response, we also tested the accumulation of PR1, a small secreted protein with antimicrobial activities. In agreement with the effect on root growth inhibition, PR1 accumulation was higher in seedlings expressing the S554/567A phosphonull mutant after BTH treatment (Figure 7D). These results, which were independent of transcription changes of the *UBQ10* promoter (Figure S6A and S6B), are also in agreement with the enhanced resistance of the S554/567A phosphonull variant to avirulent *Pst-AvrRPS4*.

## Discussion

The canonical function of the exocyst complex is exocytosis by mediating early tethering of post-Golgi vesicles before SNARE-mediated membrane fusion. Recent studies indicate that the plant exocyst can mediate polarity of the ATP-binding cassette transporter PENETRATION3 (PEN3) at the PM (Mao et al., 2016), and coordinate the transport of CASPARIAN STRIP MEMBRANE DOMAIN PROTEIN 1 (CASP1) to specific regions in the PM through the Exo70A1 subunit (Kalmbach et al., 2017). In line with these observations, Exo70B2 associates to the PM, and locates in dynamic foci and punctate compartments that concentrate at the root hair tip, a region with high secretory activity (Figure 1). Furthermore, the accumulation in BFA-compartments indicates that Exo70B2 transits through the TGN on its way to the PM or during recycling (Figure 8). This trait is, to the best of our knowledge, exclusive to Exo70B2, and suggests that it may have adopted unique features. Together, with previous results showing that Exo70B2 interacts with other exocyst subunits (Stegmann et al., 2012), our results indicate that Exo70B2 also functions as a *bona fide* exocyst subunit in secretion to the PM.

**Figure 8.**
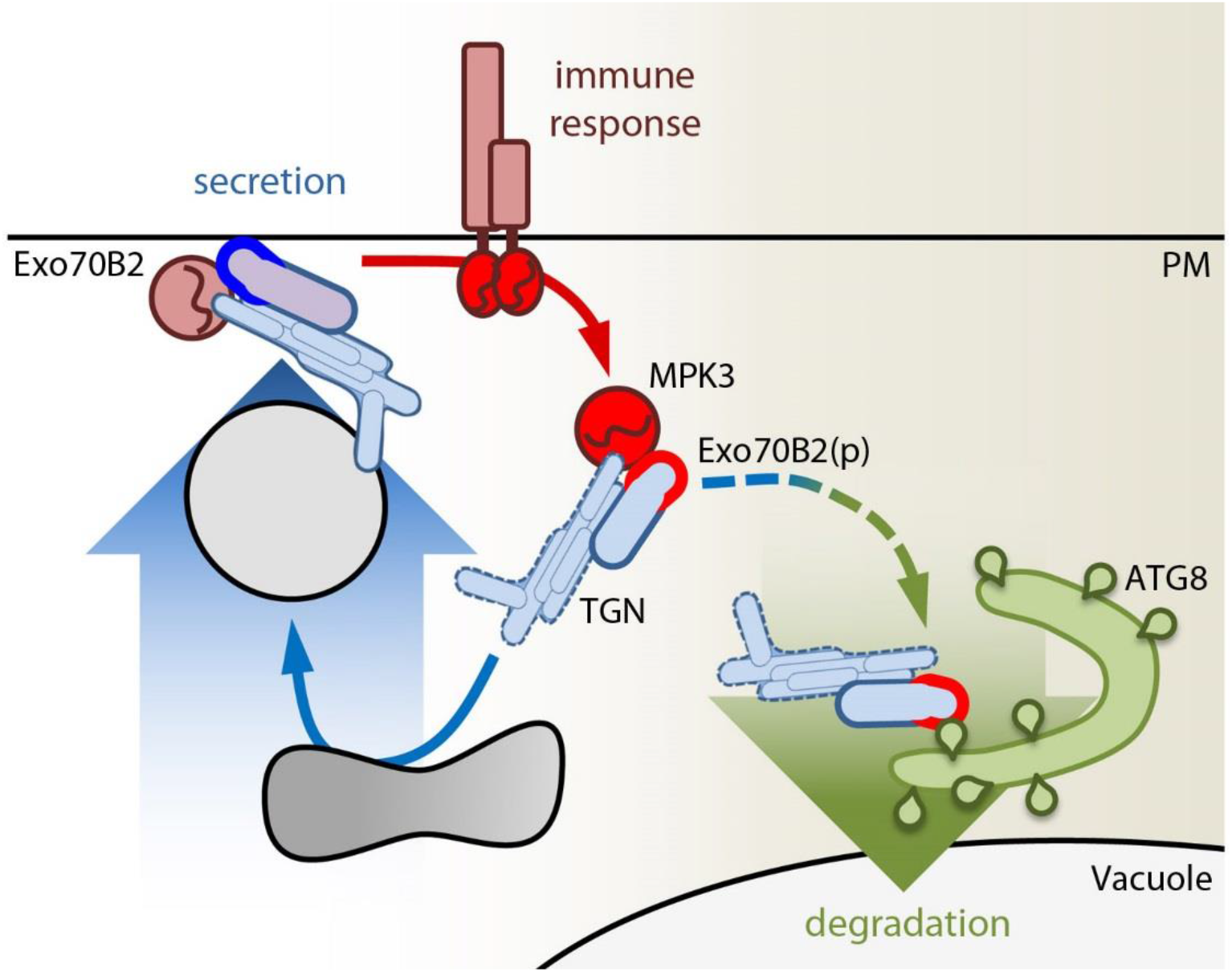
Working model for the regulation of Exo70B2 function by phosphorylation. Under basal conditions, Exo70B2 cycles between PM and the TGN and participates in secretion (blue). Activation of the immune response results in the activation of downstream signalling including MPK3, which is able to phosphorylate Exo70B2 (red). Phosphorylation reduces Exo70B2 binding to the PM. In addition Exo70B2 is degraded during the immune response, and phosphorylation may also contribute to this process by increasing interaction with ATG8. Phosphorylation thereby may mediate the gaging of the secretory activity and the cycling between membrane bound/unbound states.

The exocyst acts as a spatiotemporal regulator of secretion but many aspects pertaining the regulation of its function remain unknown. In addition to being required for the full activation of immune responses (Stegmann et al., 2012), we show that Exo70B2 interacts with and is phosphorylated by MPK3 (Figure 4), uncovering a direct link between immune signalling and the secretory machinery, which is intimately engaged in the execution and maintenance of the immune response. The identified phosphorylation sites S554 and S567 are located in the C-domain of Exo70B2, which was shown in Exo70 orthologues to harbours the surfaces mediating the interaction with phospholipids by ionic interactions (He et al., 2007a; Mei et al., 2018). The conservation of this function in plants was indirectly demonstrated with the small molecule ES2, which binds to a pocket within the C-domain and inhibits Exo70A1 function (Zhang et al., 2016; Mayers et al., 2017). Residues membrane binding in the yeast Exo70 (K605 and 608), as well as amino acids that contribute to ES2 binding to Exo70A1, are near Exo70B2 phosphorylation sites.

Moreover, our structural model suggests that phosphorylation sites S554 and S567, are placed adjoining to the last stretch of a poly-basic ridge, which agrees with previous studies that suggest an elongated interaction surface with the PM (Figure S4) (Hamburger et al., 2006; Zhao et al., 2013). Hence, introduction of negative charges by phosphorylation could regulate protein-lipid interactions (Liu et al., 2007; Zhao et al., 2013; Pleskot et al., 2015). Supporting such a scenario, the phosphonull variant S554/S567A accumulated in large patches at the root hair tip, which possesses high secretory activity, while localization was impaired in the phosphomimic variant S554/S567E (Figure 5).

This suggests that phosphorylation regulates membrane binding by introducing negative charges, dissolving the Exo70B2-phospholipid complex to undock the protein and potentially the exocyst complex. The phosphonull variant may therefore represent a constitutively active (lipid-bound) form, in which PM dissociation is impaired. In human cells, SEC3 departure from the PM occurs prior to fusion, whereas Exo70 departs just after fusion (Ahmed et al., 2018). However, it remains unclear how the departure, i.e. the disruption of the Exo70/Sec3-phospholipid complex, is mediated. Arrival at the PM also implies spatial proximity to signalling complexes, including soluble kinases, which can be recruited by other proteins (Xie et al., 2014; Zhang et al., 2015; Lin et al., 2019). Spatial proximity may result in phosphorylation of Exo70B2, and thus, its release from the membrane (Figure 8).

As part of the secretory machinery, several lines of evidence indicate that the exocyst plays a key role in plant immunity (Pecenkova et al., 2011; Stegmann et al., 2012; Ostertag et al., 2013; Stegmann et al., 2013; Fujisaki et al., 2015; Zhao et al., 2015), as well as in the innate immune system of mammals (Chien et al., 2006; Ishikawa et al., 2009). SA is a mediator of secretory activity and its accumulation, or treatment with the analogue BTH, results in a marked increase of secretion (Wang et al., 2005; Nagashima et al., 2014). Accordingly, our results show that phosphorylation of Exo70B2 is most important during conditions of high secretory activity. Plants expressing the non-phosphorylatable Exo70B2 S554/567A variant were more resistant to the avirulent *Pst-AvrRps4* strain that triggered ETI (Figure 7B). Although *exo70B2* did not show enhanced susceptibility to *Pst-AvrRPS4*, its contribution possibly is masked by a strong ETI response. In addition, Exo70B2 S554/567A plants were significantly more sensitive to BTH (Figure 7C). Enhanced sensitivity was also conferred by the WT Exo70B2, albeit significantly weaker than the S554/567A Exo70B2 variant. This observation probably reflects an altered stoichiometry of Exo70B2 to kinases such as MPK3, leading to increased levels of non-phosphorylated Exo70B2 species that result in a phosphonull-like phenotype. Moreover, the inability of the phosphonull, but not of the phosphomimetic variant, to restrict the growth inhibition response induced by BTH, underscores the importance of S554 and S567 phosphorylation.

High biosynthetic activity can pose severe strains on the secretory pathway, causing cellular stress, and activation of responses to maintain homeostasis that include autophagy (Angelos et al., 2017; Marshall and Vierstra, 2018). Subunits of the exocyst were shown to be required for autophagy activation (Bodemann et al., 2011). Two reports linked the exocyst, and more specifically its Exo70 subunits, to autophagy in plants (Kulich et al., 2013; Lin et al., 2015). Similar to observations made by Lin and colleagues (2015), we show that BTH induces the colocalization of Exo70B2 punctae with ATG8 and NBR1 (Figure 3B). In line our previous results (Stegmann et al., 2012), flg22 treatment induced Exo70B2 transport into the vacuole, where it localized to autophagosome-like punctae, and was degraded (Figure 2). Similarly, the Sec6 subunit of the exocyst also accumulated in similar compartments after BTH treatment, suggesting that other subunits follow a similar fate (Figure S1A). Supporting a conserved role in autophagy, disruption of Arabidopsis and *Brassica napus* Exo70A1 function inhibits the formation of secretory vesicles and multivesicular bodies (MVBs) at zones of pollen reception in the stigma PM (Samuel et al., 2009; Safavian et al., 2015). Similarly, for self-incompatible pollinations, secretory vesicles and MVBs are absent from the PM of stigmatic papillae, and autophagy appeared to be induced in order to redirect vesicles and MVBs to the vacuole (Safavian and Goring, 2013).

We reveal that Exo70B2 possesses two functional AIMs located in the C-domain that mediate direct binding to ATG8 (Figure 6B and 6C). The second identified AIM2 is conserved in in the majority of Exo70 homologues including Exo70E2 and Exo70B1, which colocalize with ATG8 (Kulich et al., 2013; Lin et al., 2015). Of note, AIM2 was conserved in the baker’s yeast Exo70, and able to interact with ATG8, suggesting that the interaction is conserved across kingdoms.

The two identified AIMs were near the phosphorylation sites and we show that the phosphomimic variant interacted more with ATG8 (Figure 6D and 6E). However, the S554/567A mutant variant was not significantly affected in its transport into the vacuole, suggesting that although phosphorylation may contribute to the recruitment of Exo70B2 to the vacuole via autophagy, it is dispensable, and that ubiquitination is the determining signal for degradation (Stegmann et al., 2012; Seo et al., 2016).

Together, our results indicate that various cellular pathways intersect at the exocyst complex through its Exo70B2 subunit, underscoring the importance of this protein for the coordination of secretion.

## Materials and Methods

### Plant materials and growth conditions

Arabidopsis thaliana (Col-0 accession) was used throughout this study. Seeds were surface sterilized and germinated on 1/2 Murashige Skoog media (pH 5.7) supplemented with 0.25% sucrose and 0.7% agar. Arabidopsis mutants *exo70b2-3* (GK726G07), *exo70b1-1* (GK114C03) (Stegmann et al., 2012) and *atg7-2* (GK-655B06) (Hofius et al., 2009) were previously described. All plants used for imaging were 4 day-old seedlings germinated under long day conditions (8h dark, 16h light). Protoplasts were isolated from 5-week-old plants as described before (He et al., 2007b).

### Pathogen Infection assays

For bacterial growth experiments, six-week old plants were spray inoculated with a solution of 5×10^8^ c.f.u./ml *Pseudomonas syringae* pv. *tomato* DC3000 (*Pst*) as previously described (Zipfel et al., 2004). Bacterial growth was measured 3 days after inoculation. For ETI, leaves were pressure-infiltrated with a solution of *Pst-AvrRPS4* O.D._600_ 0.001, and bacterial growth was assessed 3 days after inoculation.

### Protein purification and in vitro pull down

Recombinant MBP-Exo70B2, GST-MPK3 and GST-MPK6 were described previously (Stegmann et al., 2012). To generate recombinant His-ATG8f, coding sequence of ATG8f was cloned into pDEST17 by LR reaction. For in vitro pull-down, recombinant proteins were expressed in *Escherichia coli* after induction by 0.3mM IPTG at 28°C for 3-4h. To extract recombinant proteins, *E. coli* pellets from 50ml cultures were sonicated in 10ml column buffer (20mM Tris-HCI pH 7.4, 200mM NaCI, 1mM EDTA, 1mM DTT, 1mM PMSF, 0.5% v/v Triton X-100) and incubated at 4°C for 30 min. Extracted recombinant proteins were immobilized on either amylose or glutathione agarose resins. Thereafter, recombinant proteins of interacting candidates were added and incubated at room temperature for 1h before washing, elution and analyses by immunoblotting.

### In vitro phosphorylation assay

In vitro phosphorylation assays were performed in kinase buffer (20mM Hepes pH 7.5,15mM MgCI2, 5mM EGTA, 1mM DTT, 0.1mM ATP, 2μCi [gamma-32P]ATP) using recombinant full length MBP-Exo70B2 and active GST-MPK3, GST-MPK4 and GST-MPK11 or non-tagged MPK6 (preactivation was performed using constitutively active PcMKK5-DD) (Lee et al., 2004). Samples were incubated for 30 min at 37 °C; reactions were stopped by addition of SDS-PAGE sample buffer and separated by 10 % SDS-PAGE. Gels were stained with Coomassie Brilliant Blue and analyzed by autoradiography.

### Protein extraction and Immunoblot

To extract total proteins, 250mg fresh weight plant material was ground in liquid nitrogen to a fine powder and 2x volume buffer (10mM HEPES/KOH pH 7.5, 13.7% w/v sucrose, 5% glycerol and protease inhibitor cocktail) was added and ground to homogeneity. To obtain S10, S100 and P100 fractions, total proteins were centrifuged either for 5 min at 10k G (resulting supernatant is S10) or 30 min at 100k G (resulting supernatant and pellet are S100 and P100 fractions, respectively). Before resolving on SDS-PAGE, 1x volume of 2x Laemmli sample buffer was added to the protein and denatured at 68°C for 10min.

### Root growth inhibition assays

To monitor impact of BTH on Arabidopsis root growth, 4-day old seedlings were transferred to 1/2 Murashige Skoog media (0.25% sucrose, pH 5.7) supplemented with 100mM BTH and incubated vertically under short day conditions (8h light/16h dark). Primary root length was measured after 7 days.

### Life-imaging and Inhibitor treatment

All treatments were performed in 1/2 MS liquid media unless otherwise stated. To assay the increase in GFP-Exo70B2 fluorescence intensity, four-day old seedlings were incubated overnight in 100μM BTH. Quantification of increased fluorescence intensity (FI) signal was performed using Fiji ImageJ as described previously (McCloy et al., 2014). To visualize BFA compartments, seedlings were prestained with 5μM FM4-64 for 5min in dark followed by incubation in 50μM BFA for 45min. To quantify changes in Exo70B2 protein levels, seedlings were incubated overnight in 0.1% DMSO, 100μM BTH, 1μM ConcA, 50μM MG132 and 3h in 50μM CHX as well as indicated combinations. To visualize increased GFP-Exo70B2 and NBR1 levels, seven to ten-day old seedlings were treated for 3h with 1μM ConcA, 1h 1μM flg22 or in combination.

Subcellular localization of GFP-Exo70B2 was performed using the Zeiss LSM780n and LSM880 Airyscan laser scanning confocal microscope for high resolution images. GFP was exited using the 488 nm laser in conjunction with a 505-550 band-pass.

For Variable angle epifluorescence microscopy (VAEM), root hairs of Arabidopsis seedlings were imaged on a FEI More inverted wide field microscope (FEI, Germany) microscope using a 100 × 1.4 NA oil immersion objective. GFP was excited using a 488 nm solid-state laser diode. Fluorescence emission was collected with an EM-CCD camera with bandwidth filters ranging from 495-550 nm and fluorescence was collected with an acquisition time of 135 ms. Throughout image acquisition the microscope was operated in the ‘TIRF’ mode for vesicles in the evanescent field (the contact area between the root hair tip and the glass coverslip).

### Quantitative BiFC

Five hundred μl of protoplasts from Col-0 or *exo70b2-3* plants were co-transformed with *pSPYNE-ATG8f* (10 μg) and either *pSPYCE-Exo70B2* (13μg), *pSPYCE-Exo70B2AA* (13μg), *pSPYCE-Exo70B2DD* (13μg), *pSPYCE-Exo70B2ΔAIM1* (13μg), *pSPYCE-Exo70B2ΔAIM2* (13μg), *pSPYCE-Exo70B2ΔAIM1/2* (13μg) or empty *pSPYCE* vector (13μg). *mCherry* (7.2μg) was also co-transformed as an expression marker. Transformed protoplasts were incubated at room temperature for 14-16 h before CLSM imaging. Thirty images (with 25-40 transformed protoplast per image) were taken for each transformation event and used for quantification. The percentage of transformed protoplast (as indicated by mCherry expression) that showed YFP complementation was scored manually.

### Accession numbers

Sequence data from this article can be found in the Arabidopsis Genome Initiative under the following accession numbers: Exo70B2 (At1g07000); Exo70B1 (At5g58430); NBR1 (At4g24690); ATG8a (At4g21980); ATG8f (At4g16520); MPK3 (At3g45640); MPK4 (At4g01370); MPK6 (At2g43790); MPK11 (At1g01560).

## Acknowledgments

The authors thank Deborah Gasperini for critical reading of the manuscript and all the members of the Trujillo laboratory for fruitful discussions and valuable comments. We thank Theresa Binder for help with the generation of the yeast strain. We are grateful for materials kindly provided by Viktor Zarsky *SEC6prom:SEC6-GFP* lines, Daniel Hofius anti-NBR1, Ken Shirasu anti-HSP90, Erika Isono *pENTR-ATG8f*, and to Steingrim Svenning and Terje Johansen for *pENTR-NBR1*.

## List of Author contributions

O.K.T., C.W.L., P.M., T.K., G.F., M.Z., G.H., L.E.L., F.D. conducted experiments; T.O., W.H., J.L., M.T. designed experiments and analysed the data; M.T. wrote the paper

