## Supplemental Information for "Phosphorylation of the exocyst subunit Exo70B2 contributes to the regulation of its function"

### Supplemental Experimental Procedures

#### Yeast strains, plasmids, and growth conditions

Yeast cells were grown and manipulated according to standard procedures (Sherman, 1991, Gietz et al., 1992). The strains were cultured in synthetic complete medium containing 2% glucose as carbon source. For starvation experiments, the cells were washed and transferred to glucose-containing synthetic medium without amino acids or ammonium sulphate (Bockler and Westermann, 2014). Plasmid *pRS415-GFP-ATG8* has been described previously (Bockler and Westermann, 2014). All yeast strains are derived from BY4741 (Brachmann et al., 1998). Standard PCR-based homologous recombination was used to mark the chromosomal copy of *EXO70* with the FLAG epitope (Longtine et al., 1998).

#### Generation of transgenic lines

The cloning of Exo70B2 coding sequence into the *pENTR/D-TOPO* entry vector (Invitrogen) was described in Stegmann et al. (2012). For constitutive expression, *Exo70B2* and variants were cloned into the plant expression vector *pUBN-GFP-Dest* (Grefen et al., 2010) under control of the *UBIQUITIN10* (*UBQ10*) promoter via LR reaction (Invitrogen). Site-directed mutagenesis was performed by PCR using primers containing the desired mutation (Table S3), followed by *Dpn*I digestion. The mutation events were confirmed by sequencing. Transgenic lines were generated via *Agrobacterium tumefaciens* mediated transformation in *exo70B2-3* plants. Positive transformants were screened with 200 mg/l BASTA and confirmed via immunoblot using anti-GFP antibodies. The homozygous lines were selected based on the survival rate on full-strength MS plates supplemented with 1% sucrose and 8 g/l agar containing 50 mg/l kanamycin or 10 mg/l glufosinate.

#### Co-immunoprecipitation assays

To extract total proteins, 250mg fresh weight plant material was ground in liquid nitrogen to a fine powder and 2x volume/weight buffer (20mM HEPES, 50mM KCl, 2.5mM MgCl2, 10µM ZnCl2, 2.5mM EDTA, 5mM DTT, 0.1% Triton X-100, 10 µM 4-(2-Aminoethyl) benzenesulfonyl fluoride hydrochloride, 1mM NaF, 0.5mM Na3VO4, 15mM b-glycerophosphate, and 1% Protease Inhibitor Cocktail) was added. The lysate was cleared by centrifugation (15.000g, 10min), and the supernatant was incubated with GFP-trap beads (Chromotek) for 3 h at 4°C with gentle shaking. Beads were collected and washed five times with extraction buffer. Proteins were eluted with 2x LDS buffer (12.5mM Tris HCl, pH 6.8, 2.5% SDS, 20% glycerol, 50mMDTT) and analysed by SDS-PAGE and immunoblotting using indicated antibodies.

For coimmunoprecipitation in yeast, 30 $OD_{600}$ of cells co-expressing Exo70-FLAG and GFP-ATG8 were harvested, washed with ice-cold water, and snap-frozen in liquid nitrogen. All following steps were performed on ice or at 4°C. Native cell extracts were prepared by three rounds of 5 min beadbeating in 950 µl IP-buffer (100mM NaCl, 10% glycerol, 50mM Tris-HCl pH 8) supplemented with Roche cOmplete™ Protease Inhibitor Cocktail (Sigma-Aldrich) and 3mM PMSF. 50 µl 4% Triton X-100 in IP-buffer was added (final concentration: 0.2 %) and membranes were solubilized by rotating at 4°C for 40min. The lysate was cleared by centrifugation and incubated with GFP-nanobody resin for 4h at 4°C. The resin was washed four times with IP-buffer containing 0.1% Triton X-100 and bound proteins were eluted by addition of SDS-PAGE sample buffer and incubation at 99°C for 5min.

**Immunocytochemistry**

For ultrastructural localization, root segments were transferred in aluminum planchettes and high pressure frozen with an HPM 10 (BAL-TEC). Subsequentely the material was cryo-substituted in 0.25% glutaraldehyde (Sigma) and 0.1% uranyl acetate (Chemapol) in aceton for 2 days at -80°C using cryosubstitution equipment (FSU, BAL-TEC). This was followed by embedding in HM20 (Polysciences Europe) at -20°C.

For immunolabelling of ultrathin sections we used a polyclonal anti-gfp antibody (# 600-101-215, Rockland) detected by a rabbit anti-goat secondary antibody conjugated with 10 nm gold (#G 5527,Sigma). Sections were post-stained with uranyl acetate and lead citrate in an EM-Stain apparatus (Leica) and subsequently observed with a LIBRA 120 PLUS transmission electron microscope (Carl Zeiss Microscopy) operating at 120 kV. Images were taken with a BM-2k-120 Dual-Speed on axis SSCCD-camera (TRS).

**cDNA synthesis and Quantitative Real-time PCR**

Total RNA was prepared from adult Arabidopsis leaves using a Plant RNA Mini Kit (E.Z.N.A.® Omega bio-tek) followed by a DNaseI digestion (Thermo Fisher Scientific). For first-strand synthesis, 1 µg of total RNA was converted into cDNA with the Maxima First Strand cDNA Synthesis Kit for RT-qPCR (Thermo Fisher Scientific) according to the manufacturer's protocol. The cDNA was diluted to fixed quantities (9.5 ng per reaction of reverse transcribed total RNA).

Quantitative PCR was performed in 15 µl reaction volume, including 9.5 ng of reverse transcribed total RNA, 0.3 µM of each gene-specific primer (Table S3) and Maxima SYBR Green qPCR Master Mix 2X (Thermo Fisher Scientific). Corresponding minus reverse transcriptase and no template controls were performed with each primer pair. The qRT-PCR reaction was performed using a Bio-Rad CFX device with the following protocol: 95°C for 10 min followed by 40 cycles of 95°C for 15 sec, 60°C for 30 sec and 72°C for 30 sec, and a subsequent standard dissociation protocol to validate the presence of a unique PCR product.

In order to calculate relative transcription levels, the delta of threshold cycle (ΔCt) values were calculated by subtracting the arithmetic mean Ct values of the target Exo70B2 from the arithmetic mean Ct value of the normalizing *AtPP2A* (AT1G13320), which was obtained from the three technical replicates. The relative transcription level ($2^{-\Delta Ct}$) of each biological replicate is represented by the arithmetic mean of the three technical replicates.

**Structure modeling**

Model building was performed using the web-based SWISS-MODEL for protein structure homology modelling (Arnold et al., 2006) using the zebrafish CHIP (PDB ID code 2F42). Model quality was assessed by determining the QMEAN score: monomer Q-mean 0.647, z-score -0.55; dimer Q-mean 0.692, z-score -0.8. Images were generated using PyMol (Schrodinger, 2010).

**MS-analysis**

*In vitro* kinase assays were separated by SDS-PAGE. Coomassie G250 stained protein bands were excised, reduced with dithiotreithol (DTT), alkylated with iodoacetamide (IAA) and in-gel digested with trypsin. Peptides were desalted on in-house prepared C18 STAGE-Tips as described previously (Majovsky et al., 2014). Dried peptides were dissolved in 0.1% TFA, 5% ACN in water and injected into an EASY nLC-II liquid chromatography system (Thermo Fisher Scientific). Peptides were separated using C18 reverse phase chemistry with a column length of 10cm, a column inner diameter (ID) of 75µm, a particle size of 3µm and a 60min gradient from 5 to 40% ACN in water and a flow rate of 300nl/min. Peptides were electrosprayed on-line into an Orbitrap Velos Pro mass spectrometer (Thermo Fisher Scientific). MS/MS peptide sequencing was performed using a Top20 DDA inclusion list scan strategy to target specifically phosphorylated peptides containing the amino acid residues serine (S) and threonine (T) followed by proline (MAPK phosphorylation site motifs). Peptides were identified and phosphorylation sites mapped using the Mascot Software v. 2.5.0 (Matrix Science) linked to Proteome Discoverer v1.4. (PD, Thermo Fisher Scientific) to search the TAIR10 database (35934 sequences, 14486974 residues) and the phosphoRS module also in PD. A precursor mass error of 7ppm and a fragment ion mass error of 0.8Da was tolerated, carbamidomethylation of cysteine was set as a fixed modification, methionine oxidation and phosphorylation of S and T were set as variable modifications. A false discovery rate (FDR) was determined with the target-decoy database approach, peptide spectral matches (PSMs) with a q-value <0.05 were accepted.

Ex vivo immunoprecipitated proteins were separated and digested as above. Dried peptides were injected into an EASY nLC1000 liquid chromatography system (Thermo Fisher Scientific) and separated as above with a column length of 50cm, ID of 75µm and a particle size of 2µm using a 90 min gradient and a flow rate of 250nl/min. Peptides were electrosprayed on-line into a QExactive Plus mass spectrometer from Thermo Fisher Scientific. A Top 10 DDA inclusion list scan strategy was used to target phosphorylated peptides as above. Peptides were identified and phosphorylation sites mapped as above, tolerating a precursor ion mass error of 5 ppm and a fragment ion mass error of 0.02 Da.

### Supplemental Figures

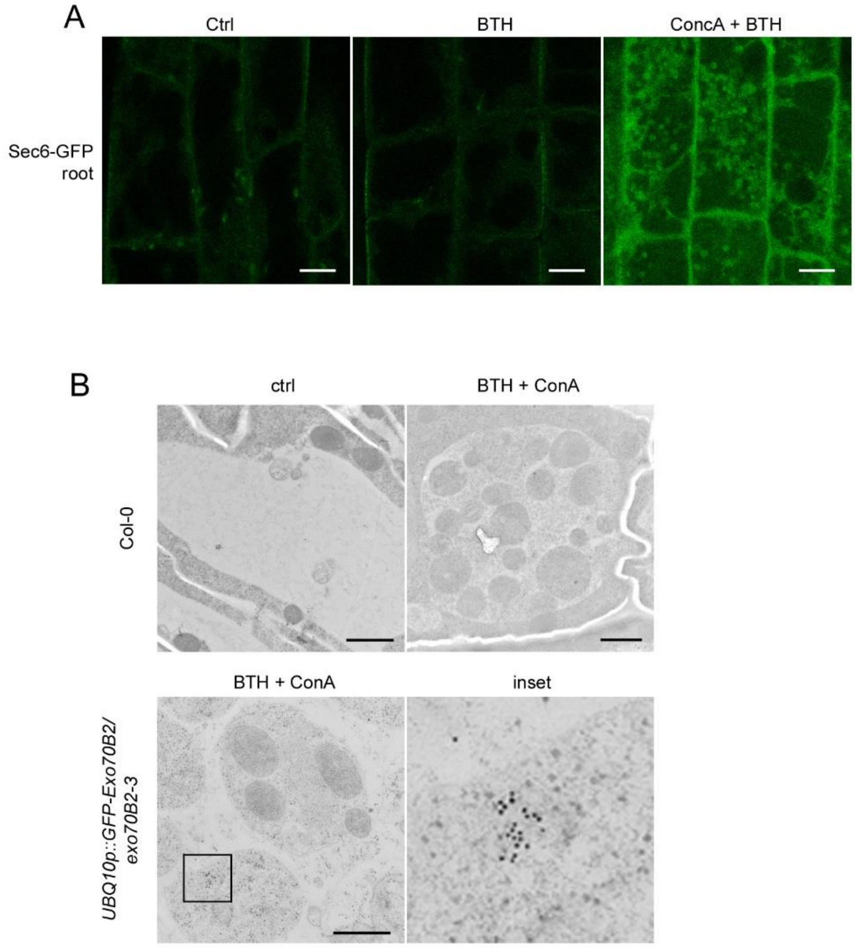

**Figure S1. Sec6-GFP is localised to autophagic-like bodies**

**(A)** Sec6-GFP is localised to autophagic-like bodies. *Sec6prom:Sec6-GFP* seedlings were treated overnight with 0.1% DMSO, 100 µM BTH or 1 µM ConcA + 100 µM BTH and analysed by CLSM. Scale bar 5 µm.

**(B)** Transmission electron microscopy pictures with GFP immuno-gold labelling. *UBQ10prom:GFP-Exo70B2/exo70b2-3* seedlings were treated overnight in 1 µM ConcA + 100 µM BTH before being subjected to cryo-fixation. Scale bars are 1 µm in the upper and 500 nm in lower panel.

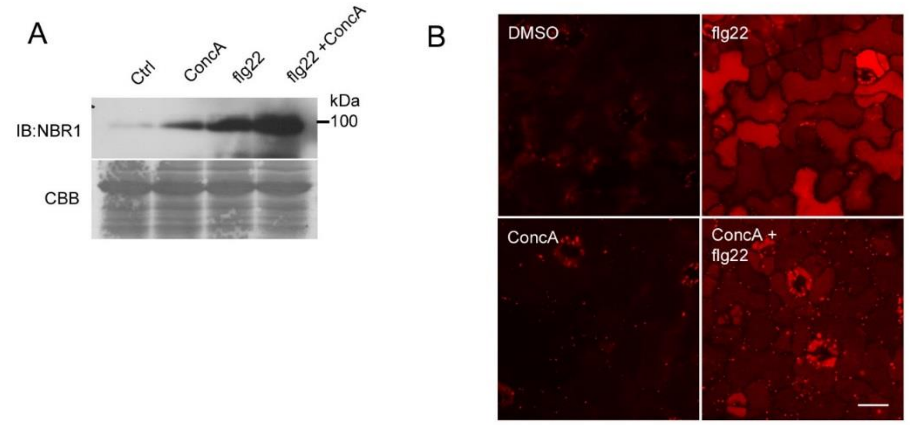

**Figure S2. The autophagic receptor NBR1 accumulates after flg22 treatment**
**(A)** Treatment with flg22 induces the accumulation of NBR1. Col-0 seedlings were grown in liquid MS for 10 days and treated for 3h with 0.1% DMSO, 1µM ConcA, 1µM flg22 or 1µM ConcA + 1µM flg22 at room temperature. Total proteins were resolved by PAGE and analysed by IB. Equal loading is shown by CBB.
**(B)** Transgenic lines carrying *UBQ10prom:RFP-NBR1* were treated overnight with 1 µM ConcA + 100µM BTH before CLSM analysis. Scale bar 20µm. (A-B) Experiments were repeated with similar results.

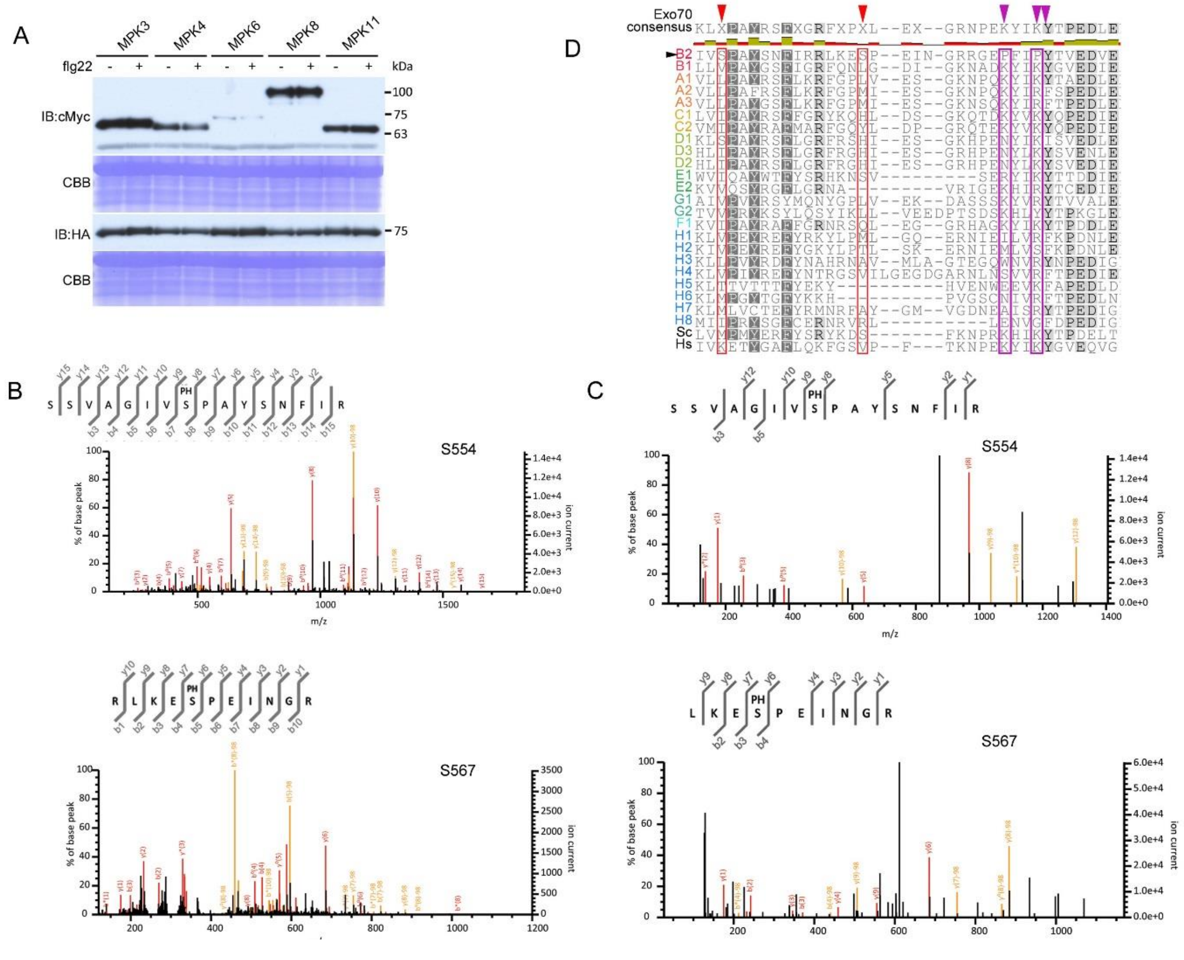

**Figure S3. Exo70B2 is phosphorylated in residues S554 and S567**

**(A)** Protein expression from BiFC experiments in which cMyc-nYFP-MPK3, cMyc-nYFP-MPK4, cMyc-nYFP-MPK6, cMyc-nYFP-MPK8 and cMyc-nYFP-MPK11 were transiently coexpressed with HA-cYFP-Exo70B2 in Arabidopsis protoplasts. Total protein samples were analysed by IB with anti-cMyc and anti-HA. Equal loading shown by coomassie brilliant blue (CBB).

**(B)** LC-MS/MS analysis spectra of two identified Exo70B2 phosphorylated tryptic peptides from in vitro phosphorylation assays with MPK3. Samples from in vitro phosphorylation reactions were resolved on an SDS-PAGE. Excised bands were consequently digested with trypsin and analysed by LC-MS/MS.

**(C)** LC-MS/MS analysis spectra of two identified Exo70B2 phosphorylated tryptic peptides from in vivo phosphorylation assay with MPK3. GFP-Exo70B2 was immunopurified from transgenic seedlings treated with flg22. Samples were resolved by SDS-PAGE. Excised bands were consequently digested with trypsin and analysed by LC-MS/MS.

**(D)** Sequence alignment of the two identified phosphorylation sites S554 and S567 in Exo70B2 with Arabidopsis Exo70 family. Also included are the sequences of the yeast (*Saccharomyces cerevisiae*, NP_012450.1) and human (NP_001138769.1) Exo70s. Phosphorylation sites are highlighted in red. Highlighted in violet are the yeast (K605 and 608) and human (K632 and K635) residues shown to influence phosphatidylinositol-binding and PM localization, and in green Exo70A1 residues (L596 and I613) influencing binding to ES2. Sequence alignment was performed by ClustalW.

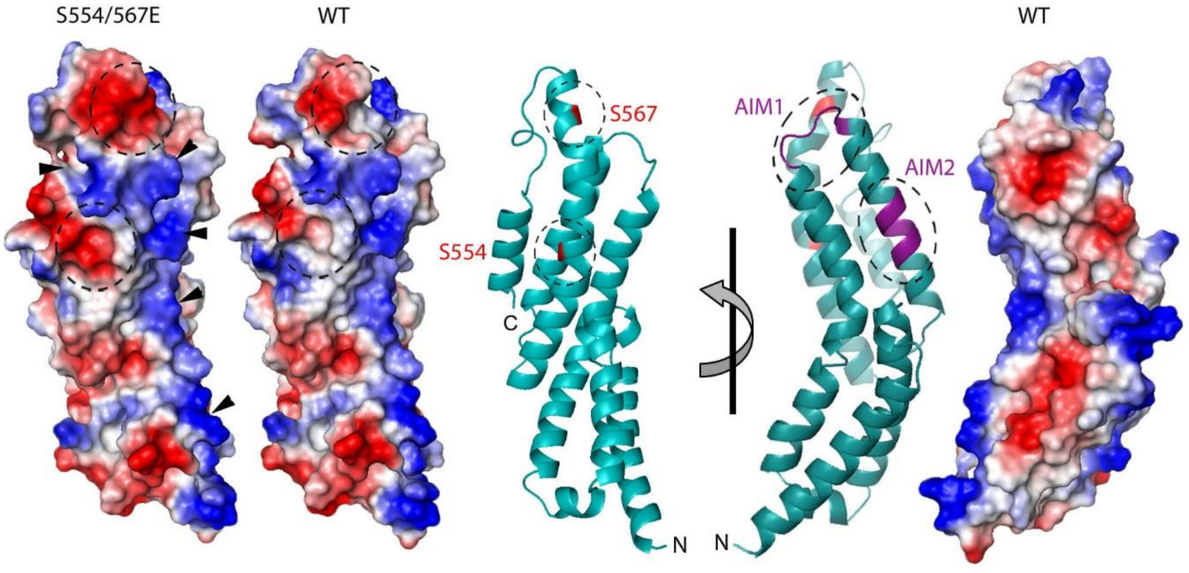

**Figure S4. Structural model of Exo70B2 C-domain**

Electrostatic surface potentials of a structural model of Exo70B2's C-domain (aa408-599) WT and the S554/567D phosphomimetic. Phosphorylation sites are highlighted in red (cartoon representation) and by dotted circles (electrostatic representation). Arrowheads indicate a predicted polybasic region. AIMs are highlighted in violet (cartoon representation).

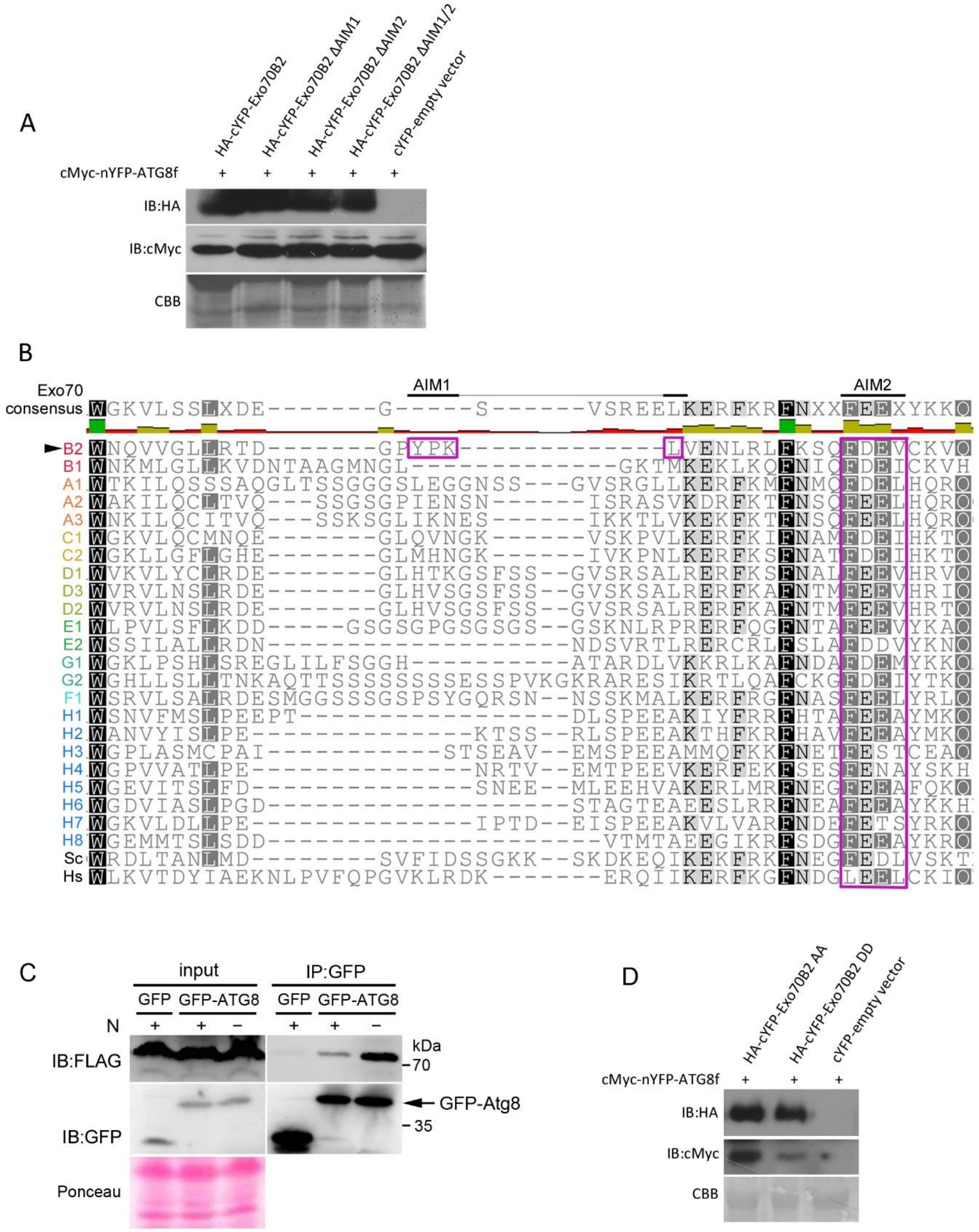

**Figure S5. Interaction of Exo70 and ATG8**

**(A)** Protein expression from BiFC experiments in which cMyc-nYFP-ATG8 was transiently coexpressed with HA-cYFP-Exo70B2 WT and mutant variants ΔAIM1, ΔAIM2, ΔAIM1/ΔAIM2 (ΔAIM1/2) or empty vector in Arabidopsis protoplasts. Total protein samples were analysed by IB with anti-cMyc and anti-HA. Equal loading shown by coomassie brilliant blue (CBB).

**(B)** Sequence alignment of the two identified AIMs in Exo70B2 with Arabidopsis Exo70 family. Also included are the sequences of the yeast (Sc, NP_012450.1) and human (Hs, NP_001138769.1) Exo70s. AIMs are highlighted in violet. Sequence alignment was performed by ClustalW.

**(C)** Yeast cells co-expressing chromosomally integrated FLAG-tagged Exo70 and either GFP* or GFP-ATG8 from a low-copy plasmid were grown to mid log phase in synthetic complete media. The cells were nitrogen-starved for 1 h, if indicated (-N). Total cell lysates were prepared and subjected to GFP immunoprecipitation followed by SDS-PAGE and immunoblot. *GFP without stop containing additional amino acids (29.7 kDa).
**(D)** Protein expression from BiFC experiments in which cMyc-nYFP-ATG8 was transiently coexpressed with HA-cYFP-Exo70B2 mutant variants S554/567A (AA), S554/567D (DD) or empty vector in Arabidopsis protoplasts. Total protein samples were analysed by IB with anti-cMyc and anti-HA. Equal loading shown by coomassie brilliant blue (CBB).

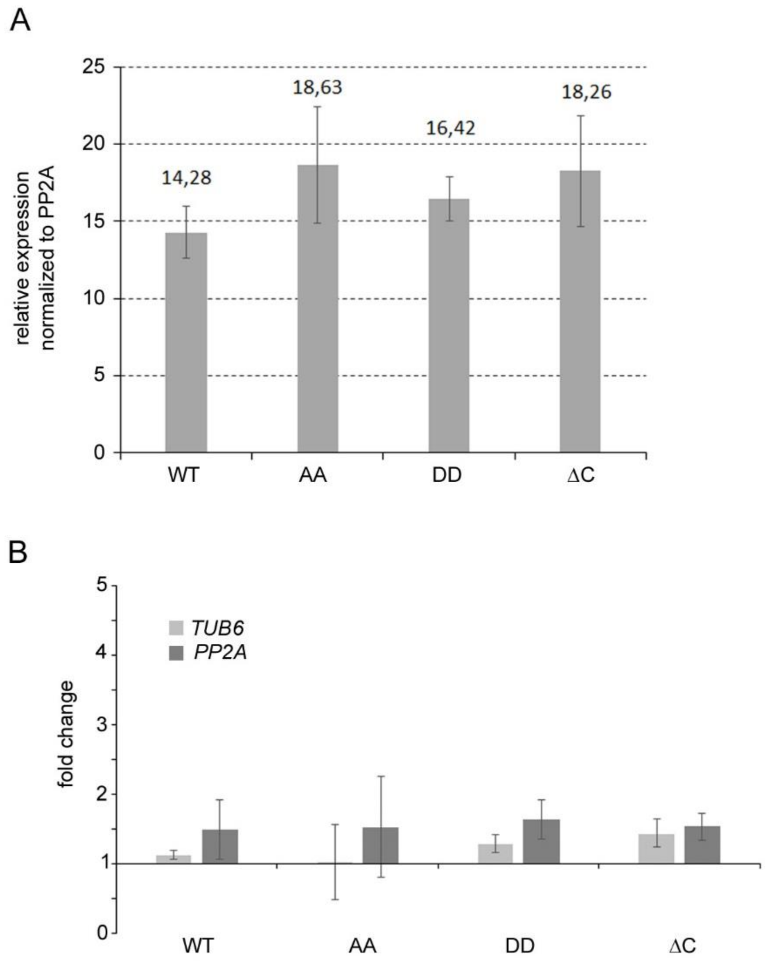

**Figure S6. Comparison of transgene transcript levels between lines and the effect of BTH treatment**

**(A)** Quantitative real-time PCR of T3 homozygous transgenic lines carrying *UBQ10prom:GFP-Exo70B2* WT and variants S554/567A (AA), S554/567D (DD) and a truncated version lacking the C-domain (ΔC). Samples were taken from adult plants and *PP2A* (AT1G13320) was used as a reference gene. Data are shown as mean +/- S.D. (n=3).
**(B)** Quantitative real-time PCR of *GFP-Exo70B2* transgene overnight treatment with 100µM BTH vs control. Expression was analysed in T3 homozygous transgenic lines carrying *UBQ10prom:GFP-Exo70B2* (WT) and variants S554/567A (AA), S554/567D (DD) and a truncated version lacking the C-domain (ΔC). *PP2A* (AT1G13320) and *TUB6* (AT5G12250) were used as reference genes to calculate relative fold change. Data are shown as mean +/- S.D. (n=3).

**Movie S1. Dynamics of Exo70B2 within the plasma membrane.** Time course of GFP-Exo70B2 detected by variable angle epifluorescence microscopy (VAEM). Vesicles at the root hair tip of five days old seedlings mounted in water. The acquisition time was 135ms per frame over 250 frames in total.

### Supplemental Tables

**Table S1. Summary of phosphorylation sites of MBP-Exo70B2 by GST-MPK3 by LC-MS/MS analysis**

| Modifications | Sample | Sequence | Mascot Ion Score | FDR Threshold | Phospho-site Probability | Charge | Number of spectra |
|---|---|---|---|---|---|---|---|
| S554 | in vivo | SSVAGIVsPAYSNFIR | 42 | 0.01 | 100 | 2 | 9 |
| S567 | in vivo | LKEsPEINGR | 24 | 0.05 | 100 | 2 | 4 |
| S554 | in vitro | SSVAGIVsPAYSNFIR | 96 | 0.01 | 100 | 2 | 172 |
| S567 | in vitro | RLKEsPEINGR | 64 | 0.01 | 100 | 3 | 8 |

**Table S2. Statistical analysis of the BTH effect on root growth**

| **One-way analysis of variance** | | | | | |
|---|---|---|---|---|---|
| P value | < 0,0001 | | | | |
| P value summary | *** | | | | |
| Are means signif. different? (P < 0.05) | Yes | | | | |
| Number of groups | 12 | | | | |
| F | 20,68 | | | | |
| R square | 0,5767 | | | | |
| Bartlett's test for equal variances | | | | | |
| Bartlett's statistic (corrected) | 14,97 | | | | |
| P value | 0,1838 | | | | |
| P value summary | ns | | | | |
| Do the variances differ signif. (P < 0.05) | No | | | | |
| ANOVA Table | SS | df | MS | | |
| Treatment (between columns) | 35,25 | 11 | 3,205 | | |
| Residual (within columns) | 25,88 | 167 | 0,155 | | |
| Total | 61,13 | 178 | | | |
| **Tukey's Multiple Comparison Test** | **Mean Diff.** | **q** | **Significant P < 0,05?** | **Summary** | **95% CI of diff** |
| Col DMSO vs Col BTH | 0,2061 | 2,009 | No | ns | -0,2772 to 0,6894 |
| Col DMSO vs exob2 DMSO | -0,08034 | 0,8147 | No | ns | -0,5450 to 0,3844 |
| Col DMSO vs exob2 BTH | 0,1354 | 1,397 | No | ns | -0,3215 to 0,5923 |
| Col DMSO vs exob1b2 DMSO | -0,2032 | 2,129 | No | ns | -0,6532 to 0,2467 |
| Col DMSO vs exob1b2 BTH | -0,09604 | 0,956 | No | ns | -0,5695 to 0,3774 |
| Col DMSO vs WT DMSO | -0,2063 | 2,012 | No | ns | -0,6896 to 0,2770 |
| Col DMSO vs WT BTH | 0,5459 | 5,068 | Yes | * | 0,03830 to 1,053 |
| Col DMSO vs AA DMSO | 0,2452 | 2,568 | No | ns | -0,2048 to 0,6951 |
| Col DMSO vs AA BTH | 1,451 | 14,45 | Yes | *** | 0,9777 to 1,925 |
| Col DMSO vs DD DMSO | -0,3418 | 3,526 | No | ns | -0,7988 to 0,1151 |

| | | | | | |
|---|---|---|---|---|---|
| Col DMSO vs DD BTH | 0,08678 | 0,895 | No | ns | -0,3701 to 0,5437 |
| Col BTH vs exob2 DMSO | -0,2864 | 2,715 | No | ns | -0,7835 to 0,2107 |
| Col BTH vs exob2 BTH | -0,07066 | 0,6799 | No | ns | -0,5605 to 0,4191 |
| Col BTH vs exob1b2 DMSO | -0,4093 | 3,991 | No | ns | -0,8926 to 0,07400 |
| Col BTH vs exob1b2 BTH | -0,3021 | 2,818 | No | ns | -0,8074 to 0,2031 |
| Col BTH vs WT DMSO | -0,4124 | 3,777 | No | ns | -0,9269 to 0,1021 |
| Col BTH vs WT BTH | 0,3398 | 2,98 | No | ns | -0,1976 to 0,8772 |
| Col BTH vs AA DMSO | 0,03911 | 0,3813 | No | ns | -0,4442 to 0,5224 |
| Col BTH vs AA BTH | 1,245 | 11,61 | Yes | *** | 0,7399 to 1,750 |
| Col BTH vs DD DMSO | -0,5479 | 5,272 | Yes | * | -1,038 to -0,05810 |
| Col BTH vs DD BTH | -0,1193 | 1,148 | No | ns | -0,6091 to 0,3705 |
| exob2 DMSO vs exob2 BTH | 0,2157 | 2,157 | No | ns | -0,2557 to 0,6872 |
| exob2 DMSO vs exob1b2 DMSO | -0,1229 | 1,246 | No | ns | -0,5876 to 0,3418 |
| exob2 DMSO vs exob1b2 BTH | -0,0157 | 0,1518 | No | ns | -0,5032 to 0,4718 |
| exob2 DMSO vs WT DMSO | -0,126 | 1,194 | No | ns | -0,6231 to 0,3711 |
| exob2 DMSO vs WT BTH | 0,6262 | 5,667 | Yes | ** | 0,1055 to 1,147 |
| exob2 DMSO vs AA DMSO | 0,3255 | 3,301 | No | ns | -0,1392 to 0,7902 |
| exob2 DMSO vs AA BTH | 1,532 | 14,81 | Yes | *** | 1,044 to 2,019 |
| exob2 DMSO vs DD DMSO | -0,2615 | 2,614 | No | ns | -0,7330 to 0,2099 |
| exob2 DMSO vs DD BTH | 0,1671 | 1,67 | No | ns | -0,3043 to 0,6386 |
| exob2 BTH vs exob1b2 DMSO | -0,3386 | 3,493 | No | ns | -0,7956 to 0,1183 |
| exob2 BTH vs exob1b2 BTH | -0,2314 | 2,272 | No | ns | -0,7115 to 0,2486 |
| exob2 BTH vs WT DMSO | -0,3417 | 3,288 | No | ns | -0,8315 to 0,1481 |
| exob2 BTH vs WT BTH | 0,4105 | 3,765 | No | ns | -0,1033 to 0,9243 |
| exob2 BTH vs AA DMSO | 0,1098 | 1,132 | No | ns | -0,3471 to 0,5667 |
| exob2 BTH vs AA BTH | 1,316 | 12,92 | Yes | *** | 0,8357 to 1,796 |
| exob2 BTH vs DD DMSO | -0,4772 | 4,849 | Yes | * | -0,9410 to -0,0134 |
| exob2 BTH vs DD BTH | -0,04862 | 0,4941 | No | ns | -0,5124 to 0,4152 |
| exob1b2 DMSO vs exob1b2 BTH | 0,1072 | 1,067 | No | ns | -0,3662 to 0,5806 |
| exob1b2 DMSO vs WT DMSO | -0,003082 | 0,0300 | No | ns | -0,4864 to 0,4802 |
| exob1b2 DMSO vs WT BTH | 0,7491 | 6,955 | Yes | *** | 0,2415 to 1,257 |
| exob1b2 DMSO vs AA DMSO | 0,4484 | 4,697 | No | ns | -0,00152 to 0,8983 |
| exob1b2 DMSO vs AA BTH | 1,654 | 16,47 | Yes | *** | 1,181 to 2,128 |
| exob1b2 DMSO vs DD DMSO | -0,1386 | 1,43 | No | ns | -0,5955 to 0,3183 |
| exob1b2 DMSO vs DD BTH | 0,29 | 2,991 | No | ns | -0,1669 to 0,7469 |
| exob1b2 BTH vs WT DMSO | -0,1103 | 1,029 | No | ns | -0,6155 to 0,3950 |
| exob1b2 BTH vs WT BTH | 0,6419 | 5,724 | Yes | ** | 0,1134 to 1,170 |
| exob1b2 BTH vs AA DMSO | 0,3412 | 3,397 | No | ns | -0,1322 to 0,8146 |
| exob1b2BTH vs AA BTH | 1,547 | 14,71 | Yes | *** | 1,051 to 2,043 |
| exob1b2 BTH vs DD DMSO | -0,2458 | 2,413 | No | ns | -0,7259 to 0,2343 |
| exob1b2 BTH vs DD BTH | 0,1828 | 1,795 | No | ns | -0,2972 to 0,6629 |
| WT DMSO vs WT BTH | 0,7522 | 6,596 | Yes | *** | 0,2148 to 1,290 |
| WT DMSO vs AA DMSO | 0,4515 | 4,402 | No | ns | -0,03181 to 0,9348 |
| WT DMSO vs AA BTH | 1,657 | 15,46 | Yes | *** | 1,152 to 2,163 |
| WT DMSO vs DD DMSO | -0,1355 | 1,304 | No | ns | -0,6253 to 0,3543 |

| | | | | | |
|---|---|---|---|---|---|
| WT DMSO vs DD BTH | 0,2931 | 2,82 | No | ns | -0,1967 to 0,7829 |
| WT BTH vs AA DMSO | -0,3007 | 2,792 | No | ns | -0,8083 to 0,2069 |
| WT BTH vs AA BTH | 0,9053 | 8,072 | Yes | *** | 0,3768 to 1,434 |
| WT BTH vs DD DMSO | -0,8877 | 8,142 | Yes | *** | -1,402 to -0,3739 |
| WT BTH vs DD BTH | -0,4591 | 4,211 | No | ns | -0,9729 to 0,05468 |
| AA DMSO vs AA BTH | 1,206 | 12 | Yes | *** | 0,7326 to 1,679 |
| AA DMSO vs DD DMSO | -0,587 | 6,054 | Yes | ** | -1,044 to -0,1301 |
| AA DMSO vs DD BTH | -0,1584 | 1,634 | No | ns | -0,6153 to 0,2985 |
| AA BTH vs DD DMSO | -1,793 | 17,6 | Yes | *** | -2,273 to -1,313 |
| AA BTH vs DD BTH | -1,364 | 13,39 | Yes | *** | -1,844 to -0,8843 |
| DD DMSO vs DD BTH | 0,4286 | 4,355 | No | ns | -0,03516 to 0,8924 |

**Table S3. List of primers used in this study**

| Primer name | Sequences (5'-3') | Purpose |
|---|---|---|
| PP2A_fwd | TAACGTGGCCAAAATGATGC | Quantitative PCR |
| PP2A_rev | GTTCTCCACAACCGCTTGGT | Quantitative PCR |
| TUB6_fwd | TGGGCTAAAGGGCATTACAC | Quantitative PCR |
| TUB6_rev | GACCTTTGGTGATGGGAAGA | Quantitative PCR |
| GFP_fwd | TGACCCTGAAGTTCATCTGC | Quantitative PCR |
| GFP_rev | GAAGTCGTGCTGCTTCATGT | Quantitative PCR |
| AIM1-APKL_fwd | ATATATGGTCTCGCTCCCAAGTTAGTAGAGAACCTAAG | AIM1 point mutation |
| AIM1-APKL_rev | ATATATGGTCTCGGGAGCCGGACCATCGGTCCTAAGCAA | AIM1 point mutation |
| AIM1-APKA_fwd | ATATATGGTCTCGCTGTAGAGAACCTAAGATTGTTCAAA | AIM1 point mutation |
| AIM1-APKA_rev | ATATATGGTCTCTACAGCCTTGGGAGCCGGACCATCGGTCCT | AIM1 point mutation |
| AIM2-ADEV_fwd | ATATATGGTCTCGCTGATGAGGTGTGTAAGGTGCAATCT | AIM2 point mutation |
| AIM2-ADEV_rev | ATATATGGTCTCATCAGCCTGTGATTTGAACAATCTTAG | AIM2 point mutation |
| AIM2-ADEA_fwd | ATATATGGTCTCGCTTGTAAGGTGCAATCTCAATGGGT | AIM2 point mutation |
| AIM2-ADEA_rev | ATATATGGTCTCACAAGCCTCATCAGCCTGTGATTTGAAC | AIM2 point mutation |
| ATG8a_fwd | ATGATCTTTGCTTGCTTGAAATT | cloning |
| ATG8a_rev | CTCAAGCAACGGTAAGAGATCCA | cloning |
